# Bright and Photostable Voltage Sensors Derived from mBaoJin

**DOI:** 10.1101/2025.05.30.657123

**Authors:** Hanbin Zhang, Shihao Zhou, Tatiana P. Kuzmicheva, Oksana M. Subach, Christiane Grimm, Minho Eom, Shulamit Baror-Sebban, Yoav Adam, Young-Gyu Yoon, Valentina Emilliani, Fedor V. Subach, Kiryl D. Piatkevich

**Affiliations:** School of Life Sciences, Zhejiang University, Hangzhou 310058, Zhejiang, China; School of Life Sciences, Westlake University, Hangzhou 310024, China; Westlake Laboratory of Life Sciences and Biomedicine, Hangzhou 310024, China; Institute of Basic Medical Sciences, Westlake Institute for Advanced Study, Hangzhou 310024, Zhejiang, China; Complex of NBICS Technologies, National Research Center “Kurchatov Institute”, Moscow 123182, Russia; Institut de la Vision, Sorbonne Université, INSERM, CNRS, F-75012 Paris, France; School of Electrical Engineering, KAIST, Daejeon, Republic of Korea; Edmond and Lily Safra Center for Brain Science, The Hebrew University of Jerusalem, Jerusalem 91904, Israel

**Author notes:** These authors contributed equally (Hanbin Zhang, Shihao Zhou).

**Keywords:** genetically encoded voltage indicators, long-term voltage imaging, in vivo voltage imaging, two-photon microscopy

## Abstract

Genetically encoded voltage indicators (GEVIs) are powerful tools for monitoring neuronal activity, but their application, particularly for long-term recordings *in vivo*, is often limited by photobleaching under the required high illumination intensities. This constraint restricts the total duration of continuous or trial-based experiments, crucial for studying processes like synaptic plasticity or circuit dynamics during behavior. Here, we introduce ElectraON and ElectraOFF, a pair of green fluorescent eFRET-based GEVIs engineered by incorporating a photostability-enhanced derivative of the bright monomeric fluorescent protein mBaoJin with Ace opsin variants. Critically, Electras demonstrate over 6-fold improved photostability compared to state-of-the-art eFRET GEVIs, pAce, and Ace-mNeon2, under one-photon illumination, while characterized by bright green fluorescence, millisecond kinetics, and good membrane localization. This enhanced stability translates to a 3-to >10-fold extension in functional recording duration, maintaining reliable spike detection in both cultured neurons *in vitro* and sparsely labeled neurons in the awake mouse cortex *in vivo*. We demonstrated sustained *in vivo* recordings exceeding 30 minutes, with instances surpassing one hour. Furthermore, Electras show functionality under scanless two-photon excitation in cultured cells. These highly photostable indicators significantly extend the temporal window for voltage imaging, broadening the scope of accessible biological questions.

## Introduction

Voltage imaging has emerged as an increasingly important tool for recording neuronal activity in model organisms *in vivo*^1–5^. Advancements in this field have primarily been driven by the development of brighter and more sensitive genetically encoded voltage indicators (GEVIs)^6–10^. Despite significant progress over the past decade, ranging from the first single-spike recordings of individual neurons in the mouse brain^8^ to population-wide imaging with single-cell, single-spike resolution, the imaging setups currently employed are relatively straightforward, utilizing wide-field microscopy with sCMOS cameras and high-power LEDs or lasers.^1,10–12^

However, a major challenge persists compared to the more established field of calcium imaging. Unlike calcium indicators, where signals arise from reporters distributed throughout the three-dimensional cell cytoplasm, voltage signals originate from GEVIs confined to the two-dimensional plasma membrane^13^. Additionally, resolving fast electrical events like single action potentials necessitates millisecond temporal resolution, requiring imaging at sampling rates from hundreds of hertz to kilohertz, which are orders of magnitude faster than those typically employed for monitoring slower calcium transients^1,10,12,13^. These combined constraints, in turn, demand the collection of a high number of photons in an extremely brief time window, necessitating illumination intensities that can range from ∼100 mW/mm² (*refs.*^4,5,9,10^) to several W/mm² *in vivo*^1,14–17^. This required photon flux is substantially higher than the intensities sufficient to resolve single-cell resolution wide-field calcium imaging, which can be over 100-fold lower^18–20^. Consequently, the required excitation power makes chromophores photobleaching the principal factor that limits the duration of high-quality voltage imaging^3,13^.

The routine recording of spikes at cellular resolution in mice using top-performing GEVIs has thus far only been extended to a few minutes under conventional microscopy^21–23^, although longer durations (30 minutes) of recording were possible in other model organisms, such as flies^10^ (**Supplementary Table 1**). This limited duration of voltage imaging is due to substantial photobleaching, leading to an inability to reliably detect single APs over time due to a drop in spike signal-to-noise ratio (SNR) in seconds, under sustained wide-field excitation^1,9,10^. Therefore, enhancing GEVI photostability is crucial for enabling longer, high-fidelity recordings, particularly given the diverse experimental subjects, preparations, and imaging systems used across different laboratories, which often necessitate adjusting imaging settings based on specific conditions^9,10,24,25^ (**Supplementary Table 1**). While advancements in GEVI engineering have often focused on optimizing parameters like brightness or sensitivity, sometimes through extensive directed evolution and laborious screening^9,10,26,27^, directly addressing the photostability bottleneck remains critical. In this study, we present ElectraON and ElectraOFF, a pair of bright green GEVIs with positive and negative polarity, respectively, designed specifically to enhance photostability. The molecular design of Electras is based on eFRET, comprising the bright and photostable mBaoJin derivative and Ace2 opsin that enables high sensitivity and millisecond kinetics. Under conventional wide-field one-photon microscopy, both Electra variants exhibited over 6-fold improved photostability in fluorescence baseline and thus, extended their functional duration for reliably reporting spike signals, increasing it from 3-to more than 10-fold compared to state-of-the-art eFRET GEVIs, including pAce and Ace-mNeon2, in both cultured cells *in vitro* and in mice *in vivo*. Additionally, we demonstrated the capability of two-photon (2P) voltage imaging with Electras in cell culture under two-photon illumination. These new indicators demonstrated outstanding photostability, underscoring their broad potential for diverse experimental demands.

## Results

### Design and optimization of Electras

Improving the fluorescent protein (FP) moiety has been a successful strategy for enhancing GEVI performance *in vivo*^10,28^. From this perspective, we sought to take advantage of two recently developed monomeric variants of StayGold, mStayGold^29^ and mBaoJin^30^, characterized by both exceptional photostability and higher intracellular brightness compared to all existing GFPs. Since most advanced GEVIs utilize GFPs as a fluorescence moiety for voltage imaging, we started with a side-by-side comparison of the established GEVIs with GFPs, including JEDI-1P^11^, JEDI-2P^27^, ASAP4e^26^, and SpikeyGi2^24^ in cultured neurons to select a suitable molecular scaffold (**Supplementary Figure 1**). For comparison, we opted to use soma-targeted variants of those GEVIs, aligning with our objective to eliminate signals from the processes, thus enhancing the extraction of somatic signals during in vivo imaging^3,31,32^ (**Supplementary Figure 1a-e**). We evaluated the performance of the GEVIs by their baseline brightness, sensitivity (ΔF/F), and SNR, which are key parameters that gauge the performance of GEVIs both *in vitro* and *in vivo*^2^. Based on the side-by-side assessment, we found that both pAce and Ace-mNeon2 outperformed other GEVIs in terms of baseline brightness as well as ΔF/F and SNR per AP when expressed in primary mouse neurons (**Supplementary Figure 1**). Consequently, we chose to proceed with the eFRET design as the prototype to engineer GEVIs using monomeric StayGold variants as the reporting moiety. In addition, this design utilizes intact GFP rather than its circular permutated version as in ASAPs^9,26^, SpikeyGi2^24^, and JEDI-2P^27^, making the engineering effort more straightforward by simply swapping one FP with another.

Correspondingly, we substituted the mNeonGreen in the soma-localized version of Ace-mNeon2 with mStayGold and mBaoJin, removing the N-terminal adaptor and adding a C-terminal c4 adaptor^33^ to mStayGold (**Figure 1a**). The resulting fusion proteins were referred to as Ace-mSG and Ace-mBJ, respectively. To generate sensors with opposite polarity, we introduced mutations into the Ace domain based on the eFRET sensor Positiron^34^ and pAce^10^ to create the positive-going pAce variant, pAce-mSG and pAce-mBJ. When we expressed these fusion proteins in HeLa cells, we observed that cells transfected with Ace-mSG and pAce-mSG exhibited no fluorescence (**Supplementary Figure 2**), while those transfected with Ace-mBJ and pAce-mBJ displayed green fluorescence localized to the membrane (**Supplementary Figure 2)**. Expression of the GEVI variants was additionally confirmed by co-expression of membrane-targeted FusionRed (FusionRed-CAAX) via P2A peptide as a reference and pAce and Ace-Neon2 as controls (**Figure 1b**). As a result, we proceeded with Ace-mBJ and pAce-mBJ to further optimize its photostability and baseline brightness.

**Figure 1.**
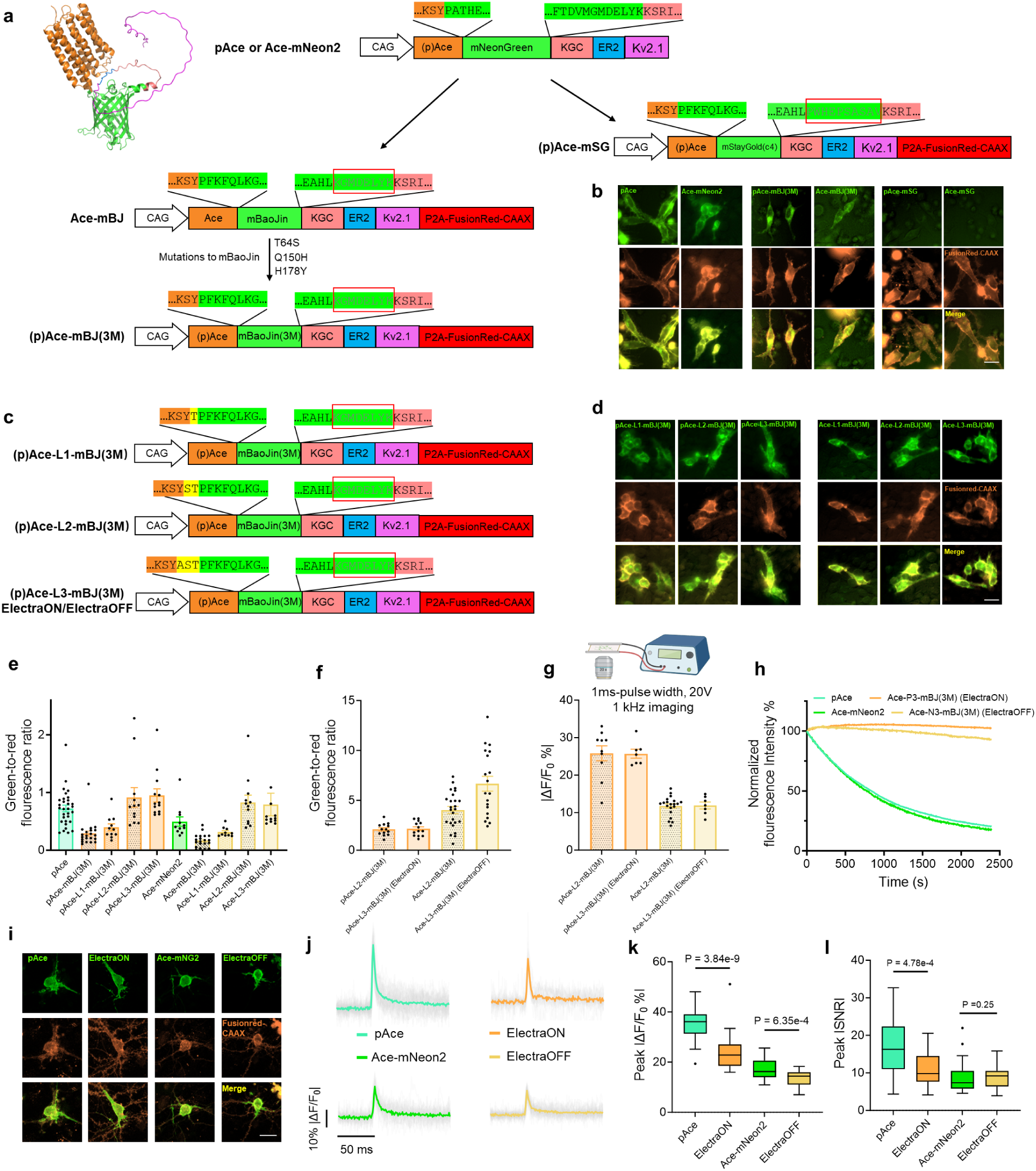
**Development and characterization of eFRET GEVIs based on mBaoJin**. **a.** Left: structure of pAce predicted by AlphaFold3(*ref.*^71^). Middle: Schematics showing the N-terminal and C-terminal adaptor sequences of mNeonGreen in pAce or Ace-mNeon2. For the generation of Ace-rhodopsin GEVIs based on mStayGold and mBaoJin, the N-terminal adaptor of mStayGold was removed and replaced mNeonGreen in pAce or Ace-mNeon2, and a c4 adaptor was added to the C terminus. The N-terminal adaptor of mBaoJin was also removed and replaced mNeonGreen in pAce or Ace-mNeon2. Three mutations were introduced to (p)Ace-mBJ to create (p)Ace-mBJ(3M), which contains mutations to enhance photostability. **b.** Wide-field images of HeLa cells expressing pAce, Ace-mNeon2, pAce-mBJ(3M), Ace-mBJ(3M), pAce-mSG, and Ace-mSG with P2A-FusionRed-CAAX post 48-h expression. **c.** Schematics showing the evolution of (p)Ace-mBJ(3M) through linker length optimization. **d.** Wide-field images of HeLa cells expressing pAce-L1-mBJ(3M), pAce-L2-mBJ(3M), pAce-L3-mBJ(3M), Ace-L1-mBJ(3M), Ace-L2-mBJ(3M), and Ace-L3-mBJ(3M) with P2A-FusionRed-CAAX under the same laser power post 48-h expression. **e.** Comparison of baseline brightness among pAce, Ace-mNeon2, and (p)Ace-mBJ(3M)-based variants in HeLa cells. The basal fluorescence of GEVI was normalized to the fluorescence of FusionRed-CAAX and reported as the green-to-red fluorescence ratio. n = 11 cells for pAce, n = 11 cells for pAce-L1-mBJ(3M), n = 12 for pAce-L2-mBJ(3M), n = 13 for pAce-L3-mBJ(3M), n = 11 for Ace-mNeon2, n = 10 for Ace-L1-mBJ(3M), n = 12 for Ace-L2-mBJ(3M), n = 12 for Ace-L3-mBJ(3M), from 2 cultures. **f.** Comparison of baseline brightness among (p)Ace-mBJ(3M)-based variants in primary hippocampal neuronal cultures, reported as the green-to-red fluorescence ratio. pAce-L3-mBJ(3M) was designated as ElectraON and Ace-L3-mBJ(3M) as ElectraOFF. n = 13 for pAce-L2-mBJ(3M), n = 13 for ElectraON, n = 25 for Ace-L2-mBJ(3M), n = 19 for ElectraOFF, from 2 cultures. **g.** Fluorescence response of GEVIs to electrical stimulation in primary hippocampal neuronal culture. n = 10 cells for pAce-L2-mBJ2, n = 7 cells for ElectraON, n = 20 cells for Ace-L2-mBJ2, n = 7 cells for ElectraOFF. **h.** Photobleaching curves of Ace-based GEVIs in HeLa cells. n = 14 cells for pAce, n = 12 cells for ElectraON, n = 12 cells for Ace-mNeon2, n = 14 cells for ElectraOFF, from 2 cultures. **i.** Representative confocal images showing the co-expression of ElectraON or ElectraOFF with the membrane reference FusionRed-CAAX in primary hippocampal neuronal culture. **j.** Averaged spike waveform generated in response to 1-AP electrical stimulation for pAce, ElectraON, Ace-mNeon2 and ElectraOFF in primary hippocampal neuronal culture. **k.** Peak ΔF/F_0_% GEVIs in response to 1-AP electrical stimulation, absolute values were shown. **l.** same as to k but for absolute SNR. n= 43 cells for pAce, 22 cells for ElectraON, 32 cells for Ace-mNeon2 and 36 cells for ElectraOFF, from two cultures. Scale bar, 25μm.

To enhance the photostability of the Ace-mBJ and pAce-mBJ, we introduced T64S, Q150H, and H178Y mutations to mBaoJin, previously identified as positions that enhance photostability in the original report^30^, and referred to this FP moiety as triple-mutated mBaoJin or mBJ(3M) for short. The resulting constructs, pAce-mBJ(3M) and Ace-mBJ(3M), generated by fusing mBJ(3M) with the pAce and Ace vectors, were both fluorescent in HeLa cell culture (**Figure 1a, b**). Further side-by-side comparison of baseline brightness in HeLa cells revealed that pAce-mBJ(3M) and Ace-mBJ(3M) were 2.5-and 1.9-fold dimmer than their mNeonGreen-based counterparts, respectively (**Figure 1e**). To increase the baseline brightness of (p)Ace-mBJ(3M), which is critical for improving SNR and, consequently, enhancing single AP detection, we extended the linker between (p)Ace and mBJ(3M) by one, two, and three aa (**Figure 1c**). Recovering two and three amino acids from the initial N-terminal adaptor of mBaoJin increased the expression-normalized baseline brightness of pAce-mBJ(3M) and pAce-mBJ(3M) by ∼3.2–3.3-and 4.7–5.0-fold, respectively, whereas the addition of one amino acid had only an incremental effect (**Figure 1d,e**). As a result, pAce-L2-mBJ(3M) and pAce-L3-mBJ(3M) were ∼1.3-fold brighter than pAce, and the brightness of Ace-L2-mBJ(3M) and Ace-L3-mBJ(3M) was 1.6-fold higher than that of Ace-mNeon2 in HeLa cells, respectively (**Figure 1d,e**). In cultured mouse neurons, pAce-L2-mBJ(3M) brightness was comparable to that of pAce-L3-mBJ(3M), while Ace-L3-mBJ(3M) was 1.65-fold brighter than Ace-L2-mBJ(3M) (**Figure 1f**). The positive-going variants exhibited almost identical ΔF/F per AP evoked by external electrical stimulation, with the value of ∼26%, while the negative-going variants showed only half of the positive variants’ amplitude of about 12% (**Figure 1g**). As a result, we chose the 3aa linker extension variants, due to their higher overall performance in neurons, for further benchmarking and validation. Correspondingly, we termed pAce-L3-mBJ(3M) as ElectraON and Ace-L3-mBJ(3M) as ElectraOFF, based on their positive and negative responses to depolarization. Altogether, these findings highlighted the direct potential of using mBaoJin variants for eFRET GEVI development, in contrast to mStayGold (**Figure 1b**), and demonstrated their matching baseline brightness compared to state-of-the-art eFRET sensors in cultured mammalian cells.

### Benchmarking of Electras in cell culture

To benchmark the performance of Electras, we systematically evaluated baseline photostability, plasma membrane localization, dynamic range, kinetics, and ΔF/F and SNR per AP using top-performing eFRET green GEVIs, pAce and Ace-mNeon2, as references. Biophysical characterization of ElectraON and ElectraOFF in cultured HEK293T cells using whole-cell patch clamp revealed their voltage-dependent response with fair linearity over −100 to +60 mV voltage steps with a dynamic range of 15.1 ± 1.8% and −20.6 ± 2.4% ΔF/F_0_, respectively (mean±SD here and throughout unless otherwise stated; **Supplementary Figure 3**), which is 3-and 2-fold smaller than dynamic range reported for pAce and Ace-mNeon2, respectively^35^. The activation and deactivation kinetics measured at room temperature matched those reported for pAce and Ace-mNeon2 measured under comparable conditions (τ_on_ = 1.11 ± 0.18 ms and τ_off_ = 0.58 ± 0.08 ms for ElectraON; τ_on_ = 1.75 ± 0.32 ms and τ_off_ = 1.14 ± 0.25 ms for ElectraOFF; **Supplementary Figure 3**; see Table S1 in ref.^10^). Both ElectraON and ElectraOFF exhibited minimal bleaching of basal fluorescence under continuous 10 mW/mm^2^ illumination intensity using wide-field microscopy, while pAce and Ace-mNeon2 lost 80% and 83% of initial fluorescence under identical conditions (**Figure 1h**). Membrane trafficking of Electras in cultured neurons was comparable to that of pAce and Ace-mNeon2, with efficient soma restriction compared to co-expression of FusionRed as a pan-membrane marker (**Figure 1i**). ElectraON and ElectraOFF exhibited 22.8% and 14.5% ΔF/F per AP, which were 37% and 11% lower compared to their counterparts, respectively (**Figure 1j,k**). However, SNR per AP for ElectraOFF was similar to that of Ace-mNeon2 (median 9.20 vs 7.36, respectively), while SNR for ElectraON was 40% lower than that of pAce (9.78 vs 16.28, respectively; **Figure 1l**). These results demonstrate the successful translation of mBaoJin properties into the engineering of photostable and functional GEVIs for reporting neuronal spikes, albeit with slightly lower sensitivity compared to top-performing eFRET GEVIs that have undergone extensive linker mutagenesis to optimize sensitivity^10^.

### Long-term recordings of Electras in cultured neurons

Building upon the superior photostability of the fluorescence baseline exhibited by ElectraON and ElectraOFF over the long term, we next investigated their utility for voltage imaging of neuronal action potentials over an extended duration (30 minutes) in cultured neurons using wide-field microscopy. We started by continuous imaging of four sensors under illumination intensity typically used for voltage imaging in cell culture. Under 5 mW/mm^2^ light power allowed us to detect single spikes with average initial SNR per AP above 4 (**Supplementary Figure 4a**). Throughout the recordings, we observed a decline in the SNR per spike across all four indicators (**Supplementary Figure 4b,c**). However, the rate of SNR decline over time was significantly slower for ElectraON and ElectraOFF compared to pAce and Ace-mNeon2 (**Supplementary Figure 4d,e**). The most pronounced reduction in SNR has been seen in Ace-mNeon2, with SNR falling below the spike detection threshold after 22 minutes, eventually merging into the noise (**Supplementary Figure 4a,c,e**). Quantitative assessment of individual neuronal spikes revealed that the initial spike SNR was 10.8 and 10.6 for pAce and Ace-mNeon2, and among the Electras, ElectraOFF had 30% higher spike SNR than ElectraON (6.09 and 4.7, respectively; **Supplementary Figure 4d,e,f**). Nonetheless, over the 30-minute recording, pAce and Ace-mNeon2 lost 50% and 60% of initial SNR values, respectively, while EelctraOFF lost only 13%, and ElectraON showed negligible reduction of SNR throughout the entire imaging (**Supplementary Figure 4f**). Furthermore, the time required for ElectraON and ElectraOFF to drop by one SNR unit was three times longer than that required for pAce and Ace-mNeon2 (**Supplementary Figure 4g**). As a result, the SNR levels of Electras could eventually surpass those of the other indicators over long-term recordings, as demonstrated by the comparison of ElectraOFF to Ace-mNeon2 (**Supplementary Figure 4**). Additionally, we confirmed that photobleaching of the fluorescence baseline was significantly less for Electras compared to pAce and Ace-mNeon2 in cultured neurons (**Supplementary Figure 4h**).

To further understand how the ultra-photostability of Electras can be exploited under different imaging parameters and applications for end users, we conducted long-term voltage recordings in cultured neurons using higher laser powers of 25 mW/mm² and 50 mW/mm², emulating the condition typically required for imaging eFRET indicators in brain tissue and *in vivo*^4,10^. Since these laser powers are five to ten times higher than the intensity sufficient for spike detection in cultured neurons^1,36^, images were acquired in trials, with each containing a 30-second continuous recording at ∼1kHz followed by a 30-second dark interval for resting and automated image data collection to minimize potential heating effects and phototoxicity by high power illumination^1,36^ (**Figure 2a**). Under 25 mW/mm² illumination, we observed an increase in spike SNR levels for all four GEVIs compared to 5 mW/mm^2^ illumination condition tested above (**Supplementary Figure 5**). The trend in SNR decline was similar to that observed at 5 mW/mm², with the best-performing neurons expressing pAce and Ace-mNeon2 showing SNRs blending into noise around sessions 19 to 20 (9.5 to 10 minutes). In contrast, the best-performing ElectraOFF continued to enable spike detection for up to 30 minutes (61 sessions) with less than a 50% drop in SNR level by the end of the recording (**Supplementary Figure 5**). Remarkably, after 7 sessions (3.5 minutes), ElectraON outperformed pAce in SNR, and after 4-5 sessions (2-2.5 minutes), the SNR level of ElectraOFF surpassed that of Ace-mNeon2.

**Figure 2.**
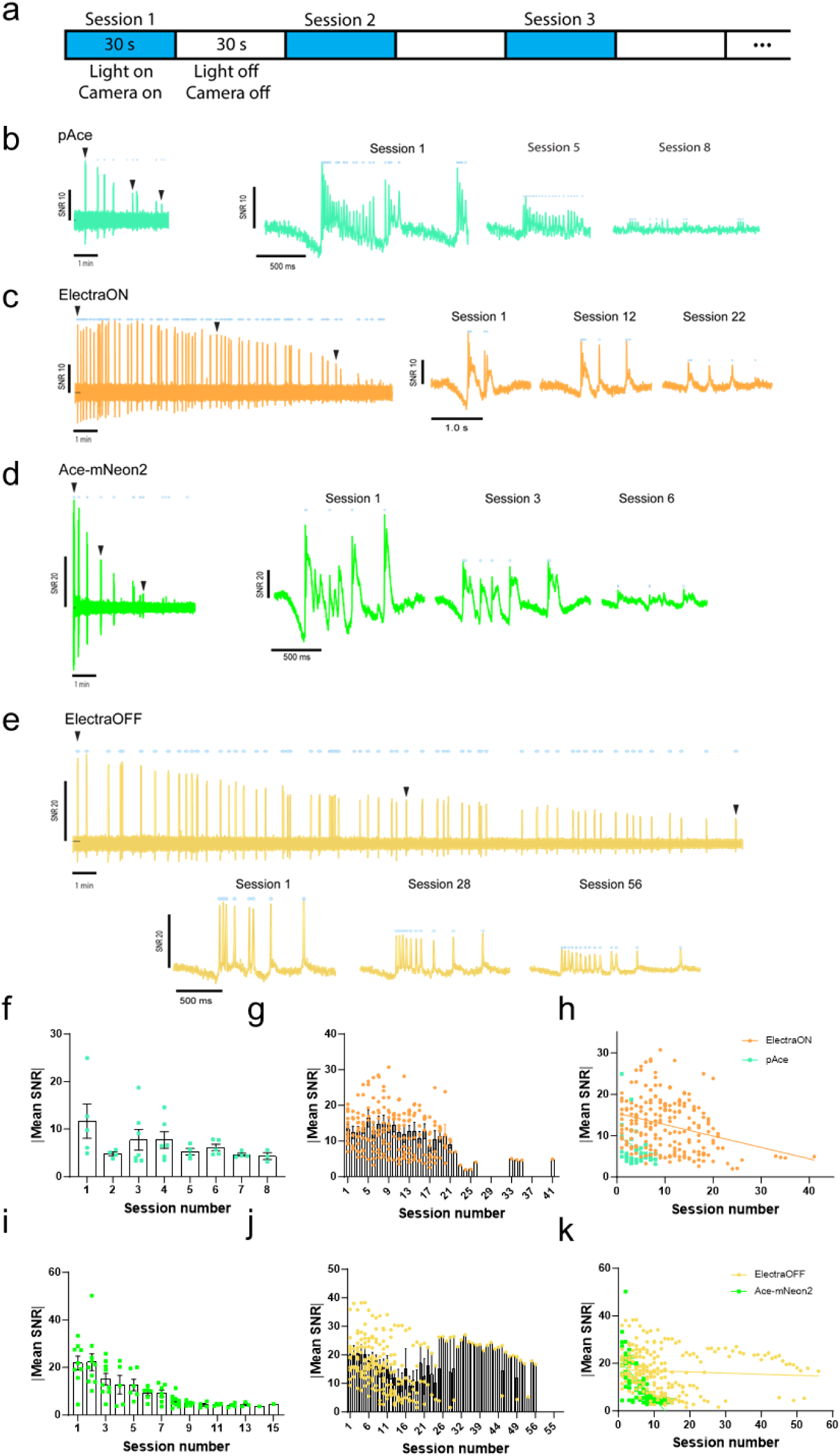
Long-term voltage imaging of cultured neurons expressing (p)Ace-based voltage sensors. **a.** Schematic diagram of the cyclic long-term voltage imaging protocol using one-photon illumination (50 mW/mm²), consisting of alternating 30-second phases: 30 seconds of neuronal voltage recording during continuous illumination, followed by a 30-second interval for automated data collection, with a total cycle duration of 60 seconds. Image acquisition rate: 996 Hz. **b.** Left panel: overview of all end-to-end 30-second traces of neuronal activity recorded from pAce-expressing neurons. Right panels: Three zoomed-in traces corresponding to the arrow-indicated regions (session1, session5 and session 8) in the overview trace, demonstrating local activity patterns and SNR level. **c-e.** Same as a but for ElectraON(c), Ace-mNeon2(d) and ElectraOFF(e). For ElectraON, zoomed-in traces corresponding to session1, session12 and session 22 were shown; For Ace-mNeon2, zoomed-in traces corresponding to session1, session3 and session 6 were shown; For ElectraOFF, zoomed-in session1, session28 and session 56 were shown. **f-g.** Absolute mean SNR value of spikes per neuron calculated for pAce(f) and ElectraON(g) across imaging sessions. n=7 neurons for pAce and n = 13 neurons for ElectraON. **h.** Linear fitting of absolute mean SNR values for pAce and ElectraON in f-g. **i-j.** Same as f-g but for Ace-mNeon2(i) and ElectraOFF(j). n=10 neurons for Ace-mNeon2 and n = 14 neurons for ElectraOFF. **k.** Linear fitting of absolute mean SNR values for Ace-mNeon2 and ElectraOFF in i-j.

Increasing the laser power to 50 mW/mm² further revealed more significant differences in SNR levels (**Figure 2b–e**) and the duration limits (**Figure 2f–k**) achieved by the four eFRET indicators. Under 50 mW/mm², neurons expressing ElectraON exhibited a spike SNR approximately three times higher (∼15 SNR units) than at 5 mW/mm² (∼5 SNR units). In contrast, SNR for pAce increased only incrementally (**Figure 2b,c,g,f**). Due to stronger bleaching under higher illumination, pAce-expressing neuron could only show detectable spikes up to a total of 4 minutes (8 sessions; **Figure 2b, f**), while ElectraON could enable spike detection up to 10.5 to 20.5 minutes (21 to 41 sessions for individual neurons; **Figure 2c, g**), with SNR level surpassed that of pAce after first imaging session (**Figure 2h**). More strikingly, the spike SNR of ElectraOFF has been scaled up to more than 3-fold (∼20 SNR unit) than that under 5 mW/mm² (∼6 SNR unit), whereas the spike SNR of Ace-mNeon2 only doubled (SNR from 10 to 20 for 5 and 50 mW/mm^2^; **Figure 2d, i**). Throughout the recording, Ace-mNeon2 was only capable of spike detection for 15 sessions (7.5 minutes; **Figure 2d, i**), whereas ElectraOFF outperformed Ace-mNeon2 in SNR within less than 2 minutes and supported recordings from 26 sessions (13 minutes) up to 56 sessions (28 minutes; **Figure 2e, j**). Altogether, these results underscore the exceptional photostability of spike fluorescence in Electras, enabling long-term voltage imaging under varied illumination conditions, thereby broadening their applicability as a proof of principle.

### Two-photon voltage imaging with Electras using parallel illumination

Two-photon excitation has been a transformative in vivo imaging tool, enabling the acquisition of high-contrast, cellular-resolution functional images in light-scattering tissues such as the mouse brain. Therefore, there have been pivotal advancements in developing two-photon voltage imaging techniques^2737^. However, the complex photocycle of eFRET voltage indicators has raised challenges for their two-photon application^38,39^, characterized by much smaller sensitivity and slower kinetics compared to those under single-photon excitation^38^. Notably, recent advancements in 2P imaging techniques have shown the possibility to improve the kinetics and sensitivity level of eFRET-opsin GEVIs by using scanless 2P strategy^40,41^, offering a path towards wider application of eFRET sensors in 2P imaging. To determine whether Electras are functional and suitable under the scanless 2P imaging modality, we expressed ElectraON and ElectraOFF in Chinese hamster ovary (CHO) cells and performed whole-cell voltage clamp recordings (**Figure 3a**). The membrane potential was stepped from-60 mV to 40 mV for 100 ms, 10 ms, and trials of 2 ms under 0.6 mW/μm^2^, which is around 40-fold less power compared to those typically used for scanning 2P (ref.^41^). During the voltage-clamp recordings, the step-response fluorescence changes of both ElectraON and ElectraOFF closely followed the applied voltage steps (**Figure 3b**), similar to observations under one-photon illumination (**Supplementary Figure 3c**). We found significant ΔF/F changes of 10% in ElectraON-expressing cells and approximately 18% in ElectraOFF, magnitudes comparable to those observed for eFRET GEVIs under scanless mode (around 20% for pAce under scanless mode, but only about 5% under scanning)^40^. The observed SNR was also in the range of 5 to 10 for both ElectraON and ElectraOFF (**Figure 3c**). In summary, these results confirm that Electras are functional under scanless 2P imaging, similar to other eFRET sensors, and suggest that their kinetics and sensitivity are significantly restored for applications under two-photon excitation.

**Figure 3.**
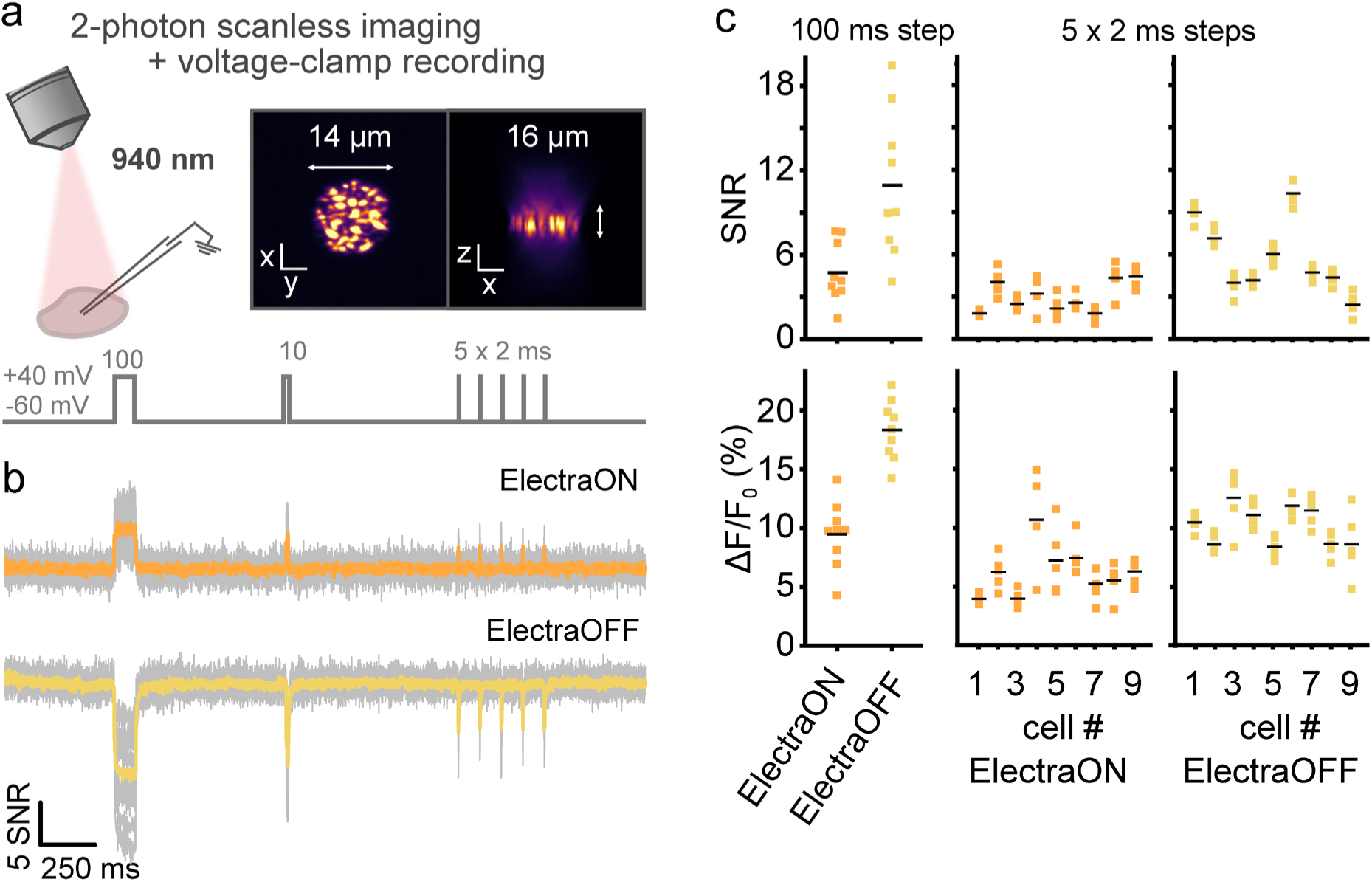
Two-photon scanless voltage imaging of Electras expressed in CHO cells. (a) Schematic of experimental recording configuration. **a.** Two-photon scanless voltage imaging was conducted using a temporally-focused holographic spot (∼14 μm lateral diameter, ∼16 μm axial resolution, 0.60 mW/μm^2^ at 940 nm) projected onto the whole cell patch-clamped cultured CHO cells expressing the Electras indicators. During recordings, the membrane potential was stepped from-60 mV to +40 mV for 100 ms, 10 ms, and five times 2 ms while fluorescence changes were imaged at 1 kHz. **b.** Averaged SNR traces for ElectraON (orange) and ElectraOFF (yellow) overlaid with single-trial individual traces (grey; n = 9 cells from one transfection each). **c.** Quantification of SNR (top) and ΔF/F (bottom) for ElectraON (orange) and OFF (yellow) for the imposed 100 ms (left) and five 2 ms depolarizations (right). Colored squares denotes individual data points and black bars denotes means.

### Longitudinal voltage imaging in mouse cortex in vivo

Encouraged by the ability of the Electra GEVIs to achieve more than 30 minutes of imaging with spike detection in neuronal culture, we proceeded with evaluating their performance *in vivo*. Given that ElectraOFF exhibited sustained but higher SNR levels than ElectraON during extended imaging in cultured neurons, we selected it for side-by-side comparison with Ace-mNeon2 *in vivo*. For imaging in awake mice, we chose conventional widefield epifluorescence one-photon microscopy (**Figure 4a**), which is both affordable and simple imaging setup, allowing direct adaptation in many laboratories for *in vivo* voltage imaging rather than more refined and complex optical systems^36,37,41,42^. ElectraOFF and Ace-mNeon2 were sparsely expressed via AAV in VIP+ interneurons of the primary visual cortex (V1), layer 2/3 (**Figure. 4b**), and imaged in head-fixed awake mice. During functional imaging, we confirmed that the indicator expression was well localized to the membrane (**Figure. 4b**). We started by setting up laser power that enabled an average signal-to-noise ratio (SNR) per action potential (AP) exceeding 4 for both voltage indicators. This threshold was selected based on established voltage imaging analysis methods using signal-level thresholding relative to noise, ensuring robust spike detection^35,43^ We observed that mean SNR per AP was comparable for Ace-mNeon2-and ElectraOFF-expressing neurons (average SNR per AP 7.44 vs. 6.26, respectively, with no statistically significant difference; **Figure. 4c**).

**Figure 4.**
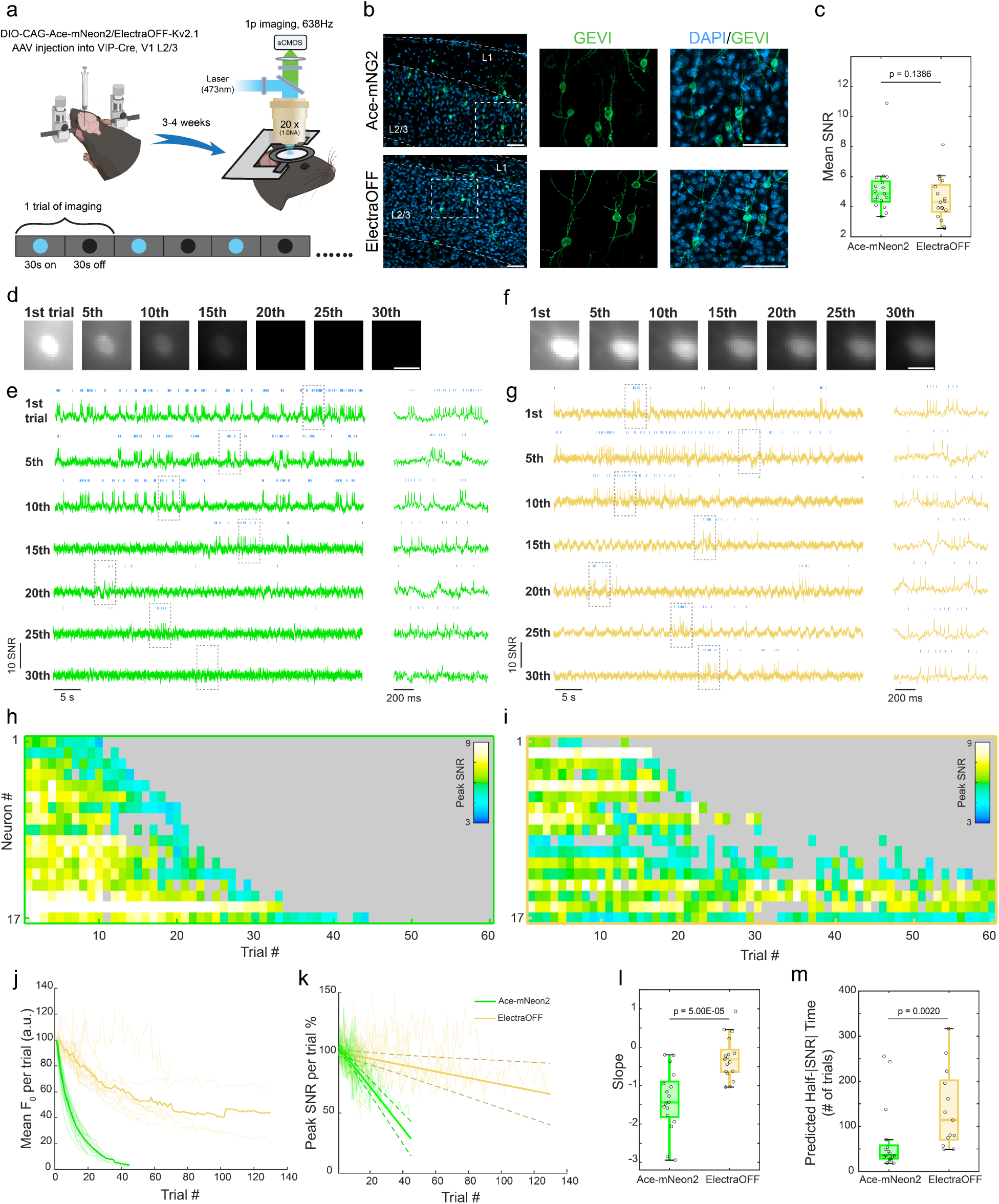
Voltage imaging of ElectraOFF expressing VIP+ interneurons in V1 using one-photon microscopy. **a.** Schematics of AAV delivery and one photon imaging experiment in awake mice. **b.** Representative confocal image showing L2/3 neurons in V1 expressing Ace-mNeon2 (top) and ElectraOFF (bottom). **c.** Mean SNR value of Ace-mNeon2 and ElectraOFF spikes recorded from VIP+ interneurons. **d.** Average projection image of an Ace-mNeon2-expressing VIP+ interneuron demonstrated for every 5th trial of recording across 30 trials. **e.** 30-sec raw fluorescence trace of the neuron shown in c demonstrated for every 5th trial of recording. Trace snippets highlighted in gray dashed box were shown on the right. **f-g.** Same as for c-d, but for ElectraOFF. Image across trials were adjusted to the identical LUT setting. **h.** Heat map depicting trial-resolved peak SNR values of spikes from recording of spontaneous activity of neurons expressing Ace-mNeon2. **i.** Same as in g but for ElectraOFF. **j.** Mean fluorescence baseline recorded over trials. Data was normalized to the fluorescence baseline recorded in the first trial. **k.** Plotting of percentage of peak SNR over trials. Data was normalized to the peak SNR obtained in the first trial. **l.** Slope quantified from the fitted line of percentage of peak SNR over trials for each neuron. **m.** Half SNR time predicted from the fitted line of percentage of peak SNR over trials for each neuron. Scale bar = 25μm. n = 17 neurons for both Ace-mNeon2 and ElectraOFF, recorded from 2 and 3 mice respectively.

We then recorded imaging trials in VIP+ interneurons, consisting of 30 seconds of continuous voltage imaging followed by a 30-second dark mode interval. These trials were repeated over an extended series of sessions until the fluorescence of the neuron blended with the background (**Fig. 4d-g**). We adapted this imaging paradigm to emulate trial-based studies in the cortex, such as neuronal plasticity^44^, skill learning^45,46^, sensory adaptation^47^, *etc.*, in which sustained, trial-by-trial monitoring is key for understanding the neural basis of behavior. In addition, this approach helps reduce heating effects and phototoxicity in the brain^48,49^, while enabling effective image data streaming, transfer, and storage. Consequently, all *in vivo* recordings were conducted in intermittent (50/50) trials rather than continuously.

Analysis of the acquired datasets revealed that ElectraOFF-expressing neurons required around 70 trials (∼35 min) to reduce their fluorescence to half of its initial baseline, which was more than six times longer than required for Ace-mNeon2-expressing neurons (specifically ∼9 trials or∼4.5 min; **Figure. 4d,e,j**). As a result of photobleaching, a pronounced decrease in SNR was observed across the Ace-mNeon2 group (**Figure. 4g, k**), while ElectraOFF showed a 4.6-fold slower decrease in SNR over time relative to Ace-mNeon2 (**Figure. 4h,k,l**), resulting in 3.2-fold more recording trials predicted (based on SNR fittings with negative-slope) before SNR declined to 50% (**Fig. 4m**). Strikingly, we observed one neuron that allow extended recording for over an hour (127 sessions, equivalent to 63 mins) in V1 (**Supplementary Figure 6**). These data underscore the exceptional photostability of ElectraOFF, extending the relative limit of voltage imaging 3–6 fold compared to Ace-mNeon2 to conduct even longer extended hour-long voltage recordings. Moreover, we found that Electras can be used to record spikes from other neuronal subtypes and brain regions, including CaMKII+ neurons and those from the hippocampus (**Supplementary Figure 7**).

## Discussion

A critical bottleneck limiting the application of voltage imaging, especially for longitudinal studies in vivo, has been the rapid photobleaching of GEVIs under the intense illumination typically used to record fast neural dynamics. The achievable recording durations, often just minutes with top-performing GEVIs in mice (**Supplementary Table 1**), are insufficient for many neuroscience applications demanding extended observations. Examples include monitoring synaptic plasticity phenomena like long-term potentiation (LTP) and long-term depression (LTD), which evolve over tens of minutes to hours^50,51^, tracking potentially slow shifts in firing patterns (e.g., burst-firing transitions) associated with learning^16^ or chronic mental disorders such as depression^52,53^. While voltage imaging has provided valuable snapshots of neural activity during behavior^1,4,10,16^, extending the recording window is essential for dissecting the mechanisms underlying longer timescale processes.

In this study, we addressed this limitation by developing ElectraON and ElectraOFF, leveraging the exceptional photostability of the mBaoJin variants within an eFRET GEVI framework. By directly substituting mNeonGreen in the well-characterized (p)Ace scaffold with a photostability-enhanced mBaoJin variant mBJ(3M), we achieved a dramatic improvement in photostability (>6-fold reduction in baseline bleaching) with minimal optimization effort. Although this came with a decrease in initial single-AP sensitivity (ΔF/F) compared to the highly optimized pAce/Ace-mNeon2 sensors, the practical consequence was a significantly slower decay of spike SNR during prolonged imaging both in cell culture and in vivo in mice. In cultured neurons, Electras enabled reliable spike detection for over 30 minutes under moderate illumination, far exceeding the functional lifetime of the used references. Under high illumination intensities mimicking *in vivo* conditions, Electras not only lasted 3-to >10-fold longer but their SNR levels surpassed those of pAce and Ace-mNeon2 within minutes, highlighting their advantage for demanding, long-duration experiments. Crucially, this improved photostability translated robustly to the *in vivo* setting. ElectraOFF enabled sustained wide-field 1P imaging of VIP+ interneurons in awake mice for >30 minutes, with quantifiable 3-to 6-fold improvements in baseline stability and functional SNR duration compared to Ace-mNeon2. In the parallel study, Wang et al. applied ElectraOFF for robust voltage imaging of hippocampal CA1 pyramidal neurons, visual cortical parvalbumin interneurons, and striatal cholinergic interneurons over tens of minutes, occasionally up to eighty minutes, in behaving mice^54^. Altogether, these results established ElectraOFF as an enhanced green GEVIs, allowing the long-term in vivo voltage imaging of various neuronal cell types, thus representing a substantial advancement for the field.

Furthermore, our demonstration of ElectraON and ElectraOFF functionality under scanless 2P illumination suggests their potential applicability for deep-tissue imaging, where their photostability could be particularly beneficial, although further *in vivo* 2P characterization will be required. The results underscore the power of optimizing the FP component for GEVI development, showing that incorporating intrinsically more photostable FPs like mBaoJin derivatives can yield significant gains in functional imaging duration.

While Electras offer a major advantage in photostability, the slightly lower initial sensitivity might be a consideration for applications detecting extremely small subthreshold events or requiring maximal SNR in very short recordings. However, for the many experiments limited by photobleaching – including studies of learning, plasticity, network states, and disease models operating on timescales of many minutes to hours – the extended functional duration provided by Electras represents a significant enabling technology. Future iterations could potentially combine the mBaoJin scaffold with further protein engineering to optimize sensitivity while retaining high photostability. Our results indicate that mBaoJin may be a more suitable FP for sensor development than mStayGold, which appears non-fluorescent when fused with Ace (**Supplementary Figure 2**). This suitability could be due to mBaoJin’s enhanced folding and maturation time, taking only approximately 7.6 minutes^55^. Furthermore, a recent study by Sekhon et al. also identified mBaoJin as the most suitable FP among all monomeric versions of StayGold for biosensor development^56^.

In the parallel study, Cao et al. explored an alternative eFRET GEVI design by inserting the enhanced mBaoJin variant into Ace2 rhodopsin’s first extracellular loop^57^. The obtained GEVI, named Vega, combines high brightness, sensitivity (-33% ΔF/F per 100 mV), fast response (1.4 ms), with enhanced photostability, enabling continuous, high-fidelity voltage imaging in neurons and pancreatic islets. Vega enabled stable, long-term recordings in acute brain slices, pancreatic islets, and awake mice, revealing detailed neuronal firing patterns and cellular activity dynamics with minimal photobleaching.

In summary, ElectraON and ElectraOFF provide a valuable combination of brightness, speed, and dramatically improved photostability. They significantly extend the achievable duration for reliable voltage imaging in vitro and in vivo, expanding the range of biological questions accessible with optical electrophysiology.

## Methods

### Molecular Cloning

The genes of pAce, Ace-mNeon2, ASAP4e, SpikeyGi2, JEDI-1P, and JEDI-2P were *de novo* synthesized by Synbiob Gene Technology Co., China, based on the sequence reported in the original publications. To clone the pAAV-CAG-ElectraON-P2A-FusionRed-CAAX and pAAV-CAG-ElectraOFF-P2A-FusionRed-CAAX plasmids, the genes were PCR amplified with KpnI/AgeI flanking sites and swapped with the StayGold-P2A-mCherry gene in pAAV-CAG-StayGold-P2A-mCherry (WeKwikGene plasmid#0000293). Synthetic DNA oligonucleotides used for cloning were synthesized by Tsingke Biotechnology Co. or Zhejiang Youkang Biological Technology Co., China. ApexHF HS DNA Polymerase FS Master Mix from Accurate Biotechnology (Hunan Co., China) or PrimeStar Max master mix (Takara, Japan) was used for high-fidelity PCR amplifications. Restriction endonucleases were purchased from New England BioLabs and used according to the manufacturer’s protocols. DNA ligations were performed using an OK Clon DNA Ligation kit II from Accurate Biotechnology (Hunan Co., China). The ligation products were chemically transformed into the TOP10 *E. coli* strain (Biomed, China) and cultured according to the standard protocols. Sequencing of bacterial colonies and purified plasmids was performed using Sanger sequencing at Zhejiang Youkang Biological Technology Co., China. Small-scale isolation of plasmid DNA was performed with commercially available Mini-Prep kits; large-scale DNA plasmid purification was performed with Midi-Prep kits (QIAGEN, Germany). The mammalian plasmids, including fully annotated and verified sequences, generated in this study are available from the WeKwikGene plasmid repository at Westlake Laboratory (https://wekwikgene.wllsb.edu.cn/).

### Characterization of ElectraON and ElectraOFF variants in cultured mammalian cells

HEK293FT (Invitrogen, USA) and HeLa (ATCC CCL-2) were authenticated by the manufacturer using short tandem repeat profiling, reauthenticated in our laboratory by inspecting stereotypical morphological features under a wide-field microscope, and tested negative for Mycoplasma contamination to standard levels of stringency. Authentication by morphology was performed every time before transient transfection. HEK293FT and HeLa cells were cultured in DMEM supplemented with 10% fetal bovine serum and 1% penicillin/streptomycin and seeded in a 24-well glass-bottom plate (lot#P24-0-N, Cellvis, USA) or glass-bottom dish (MatTek, USA) after Matrigel coating (lot#356235, BD Biosciences, USA) conducted by 30 min incubation at 37°C. Cells were transfected at 80–90% confluency using Hieff Trans Liposomal Transfection Reagent (lot#40802ES02, Yeasen Biotechnology, China) according to the manufacturer’s protocol and imaged 24–48 h after transfection. Cells were imaged using a Nikon Ti2-E widefield fluorescence microscope equipped with a Spectra III Light Engine (LumenCore, USA), an ORCA-Flash 4.0 V3 sCMOS camera (Hamamatsu, Japan), ×10 NA 0.45 and ×20 NA 0.75 objective lenses (Nikon, Japan) controlled by NIS Elements AR 5.21.00 (Nikon, Japan).

To measure the brightness of the Electra variants in live cells under a widefield microscope, HeLa cells were transfected with the pAAV-CAG-Electras-P2A-FusionRed-CAAX plasmids and were imaged in FITC (excitation 475/28 nm from Spectra III LumenCor; emission 594/40 nm) and TRITC (excitation 555/28 nm from Spectra III LumenCor; emission 535/46 nm) channels. To obtain statistically significant datasets, we performed two independent transfections, and regions of interest (ROIs) were determined using the auto-detect function of NIS Elements software, limiting the ROI area to 50 μm^2^ as a minimal size of HeLa cells. The mean fluorescence intensity in FITC and TRITC channels for ROIs was extracted and the FITC:TRITC ratios were calculated after background subtraction for each channel, which was used for the comparison of intracellular brightness under corresponding imaging conditions. The data were excluded from the analysis if cells were dead or out of focus.

To measure photostability in live HeLa cells, HeLa cells were transfected as described above and imaged under continuous FITC widefield excitation. The 20x NA0.75 objective lens (Nikon) and Spectra III LumenCor were used and set at 20% power. The illumination power at the focal spot was 10 mW/mm^2^. Photobleaching curves were calculated for each cell individually and reported as a mean photobleaching curve for each protein (averaged from all individual curves). Cells that detached or died during photobleaching experiments were excluded from data analysis.

Dissociated neuronal cultures were prepared in-house from the C57BL/6 mice (supplied by the animal facility of Westlake University) at postnatal days 0–1, regardless of sex, following a previously reported protocol^58^ and in strict accordance with the Westlake University Animal Care Guidelines approved by the Institutional Animal Care and Use Committee (IACUC) of Westlake University, Hangzhou, China, under animal protocol 19-044-KP. Briefly, the hippocampi from pups were quickly isolated under a stereomicroscope and washed with ice-cold DMEM/High glucose (G4510, Servicebio, China) and incubated with 0.25% trypsin (25200056, GIBCO, USA) for 12 min at 37°C while gently shaking every 4 min to facilitate digestion. The digestion was terminated by adding DMEM/High glucose supplemented with 10% FBS preheated at 37°C. The digested tissue was then gently dissociated using Pasteur pipettes with 10 µl tip. The cell suspension was pelleted and resuspended with fresh medium, then filtered through a 40-μm nylon strainer. The dissociated neurons were plated at a density of 150,000 per well on pre-coated coverslips (1254580, ThermoFisher Scientific™, USA) with Matrigel (356235, BD Biosciences, USA) in Advanced MEM containing 10% FBS-ES as seeding medium. After 12 h incubation at 37°C/5% CO_2_ for cell adhesion, half of the medium was replaced by Neurobasal medium (21103049, GIBCO, UK) supplemented with 1% GlutaMax (35050061, GIBCO, UK) and 2% B27 (17504044, GIBCO, UK). AraC (4 μM, C6645, Sigma, USA) was added when glia cell density was 70–80%.

To quantify SNR spike induced by field stimulation using a self-built electrical system described earlier in live primary mouse hippocampal neurons^59^. Cultured neurons were transfected with 500 ng of pAAV-CAG-Electras-P2A-FusionRed-CAAX plasmids per well using the calcium phosphate transfection method or transduced using rAAV2/9-CaMKII-Electras viruses (Shanghai Sunbio Medical Biotechnology, China). In brief, hippocampal neurons were transfected at DIV (days in vitro) 4–5 using the calcium phosphate transfection kit (lot#K278001, Thermo, USA) according to the previously described protocol^60^. To obtain a high density of neurons expressing GEVIs, transduction was performed with ∼1 µl rAAV2/9 viruses (>10^12^ vector genome (v.g.) per ml) per well, in 1 ml culture medium at DIV 4–5. Neurons were imaged at DIV 12–18 using the same imaging setup as described above for HeLa cells. To increase the neuronal activity in room temperature (22℃)during imaging, neurons were treated with a GABA_A_ receptor antagonist bicuculline (BIC, 30 μM, HY-N0219 from MCE) and a potassium channel antagonist 4-aminopyridine (4AP, 100 μM, HY-B0604 from MCE). Mean fluorescence intensity was calculated for neuronal soma only. The culture was excluded from the analysis if cells died after transfection or if the culture was contaminated with bacteria or yeast.

### Data processing and analysis for long-term intermittent imaging in cultured neurons

The analysis of time-series voltage imaging data was performed using custom MATLAB (MathWorks) scripts, adapting and implementing procedures based on established methodologies^61^.The processing pipeline involved motion correction, PCA-based denoising, region of interest (ROI) definition, trace extraction, spike detection, and signal-to-noise ratio (SNR) calculation.

*(1) Motion Correction and Image Preprocessing:* Raw image sequences (.tif files) were first loaded as image stacks (ImStack). To correct for sample motion primarily occurring in the x-y plane, a 2D motion correction algorithm was applied. A reference template frame (Template) was generated by calculating the temporal mean across the entire image stack. Motion shifts (Shift) between each frame in the stack and the template were estimated using a phase-correlation-based method (motion_est function). This involved calculating the normalized cross-correlation between the spatial gradients (approximated using complex representation (I(x) - I(x-1)) + 1i*(I(y) - I(y-1))) of the template and each frame within a defined region (typically the full frame, ROIPos). To avoid edge artifacts influencing the correlation, a mask (Mask) was applied, excluding pixels near the border (MaxShift = 4 pixels). Peak fitting on the cross-correlation map was used to determine the x and y shifts for each frame with subpixel accuracy (XShiftFine, YShiftFine). The estimated shifts were then used to correct the original image stack (ImStack) using 2D linear interpolation (motion_crt function using interp2), generating a motion-corrected stack (ImStackCrt). Edge pixels potentially affected by the maximum interpolation shift were masked out. Subsequently, adjacent frames in the corrected stack were averaged (ImStackCrt(:,:, 2:end) + ImStackCrt(:,:, 1:end-1)) to reduce potential artifacts arising from bidirectional scanning and improve the signal-to-noise ratio. The motion-corrected, frame-averaged stack was saved (_corrected.mat).
*(2) PCA-Based Denoising:* To reduce noise uncorrelated across pixels, Principal Component Analysis (PCA) based denoising was applied. The motion-corrected stack (ImStackCrt) was spatially downsampled by block averaging using a block size (BNxy = 8×8 pixels) to create ImPCA. Singular Value Decomposition (svds) was performed on this temporally concatenated, downsampled data (ImPCA) to extract the principal components (PCs) of the dominant spatiotemporal patterns, capturing potential correlated noise or widespread background fluctuations (noise_pca_extract function). The top principal component(s) (NoisePCA.Patterns, NoisePCA.Trace, typically NoiseCrtN=1) were identified as representing noise patterns. A denoised image stack (ImStack) was then generated by removing the contribution of these noise components from the full-resolution motion-corrected stack (ImStackCrt) via linear regression (noise_pca_crt function). Specifically, the projection of each noise component’s temporal trace (NoisePCA.Trace) onto the data was calculated, and this scaled noise pattern was subtracted from the original data.
*(3) ROI Definition and Trace Extraction*: Regions of Interest (ROIs) corresponding to individual neurons were manually defined using the imfreehand tool in MATLAB on a representative image (e.g., the motion correction Template or a standard deviation projection image). The pixel indices for each manually drawn ROI were saved (ROI_…mat). For each FOV, these predefined ROIs (ROIs) were loaded. To extract the fluorescence trace for each neuron, the denoised image stack (ImStack) was processed. For a given ROI (Neuron(ii).ROI), the raw fluorescence trace (RawTrace) was calculated as the temporal evolution of the mean pixel intensity within the ROI mask (SomaIdx) across all frames (extract_roi function).
*(4) Trace Processing, Spike Detection, and SNR Calculation:* For each extracted raw ROI trace, several processing steps were performed (spike_extract function). Local Background Subtraction: A local background region was defined for each ROI, encompassing pixels within a defined radius (Param.CellEnvSize = 15 pixels) surrounding the ROI’s bounding box, but excluding pixels belonging to any defined ROI (BkgIdx derived from ROITot). The mean fluorescence trace of this local background region (Bkg) was calculated and subtracted from the ROI’s raw trace to yield the background-corrected trace (Trace = RawTrace - Bkg). Filtering: The background-corrected trace was high-pass filtered using a third-order Butterworth filter (butter, filtfilt) with a cutoff frequency (Param.CutOffFreq = 1 Hz) to remove slow baseline drifts (highpass_video_filt function), resulting in FiltTrace. Noise Estimation: The standard deviation of the noise (noiseStd or NoiseAmp1) was estimated from the filtered trace (FiltTrace). Assuming negative-going voltage spikes (Param.SpikePolarity =-1, though the code uses +1 which might need checking based on the indicator used), the noise level was estimated from the segments of the trace presumed not to contain spikes (*e.g.*, using parts of the trace excluding spike windows, as implemented in calculateSNR, or potentially using the positive half as mentioned in the reference paper). Spike Detection: Putative spike times (SpikeIdx) were identified using a spike detection algorithm (spike_denoise function, details inferred partially from parameters and the reference paper). This process typically involved i) initial detection of peaks in the filtered trace exceeding a signal-to-noise ratio threshold (e.g., Param.SNRList(1) = 4); ii) generation of an average spike waveform template by aligning and averaging trace segments around initially detected spike times (Param.SpikeTemplateLength); iii) refinement of spike times, potentially using template matching or matched filtering techniques, followed by adaptive thresholding to obtain the final set of spike indices (SpikeInfo.SpikeIdx). The SpikeInfo structure stored the detected spike times and related trace information. Raw traces (SpikeInfo.RawTrace) were also stored, even for ROIs classified as non-spiking. SNR Calculation: The Signal-to-Noise Ratio (SNR) for each neuron was calculated (calculateSNR function). The average spike amplitude (avgAmp) was determined by measuring the peak amplitude of detected spikes in the filtered trace relative to the local baseline preceding each spike. The noise level (noiseStd) was estimated as described above, carefully excluding windows around detected spikes. The final reported SNR for a neuron was the ratio avgAmp / noiseStd. These SNR values were compiled across all neurons and FOVs for analysis.

### Data processing and analysis of 30-min long-term voltage recording from cultured neurons

To process the longitudinal voltage imaging dataset recorded at 1000 Hz, 30 minutes, the.nd2 files were first converted into an HDF5 format to enable efficient random access from disk without requiring full RAM loading. Motion correction was performed using NoRMCorre^62^ integrated in the pipeline of Volpy, applying a non-rigid motion correction algorithm. Following motion correction, neural activity and corresponding spike events were extracted using Volpy, an automated and scalable analysis pipeline optimized for voltage imaging datasets. Given that only a single neuron was acquired per FOV, for precise region-of-interest (ROI) annotations, we used manually segmented ROIs from ImageJ instead of automatic annotation. Signal polarity of processing was determined according to the positive or negative fluorescence change of the GEVIs. The trace was then high-pass filtered (1/3Hz) to correct for photobleaching, followed by spike detection using a whitened match filter and adaptive thresholding embedded in the algorithm.

Subsequent analyses were performed based on the neural activity and spike data extracted by Volpy. To quantify signal-to-noise ratio (SNR) over time, the SNR trace was computed using the following equation:

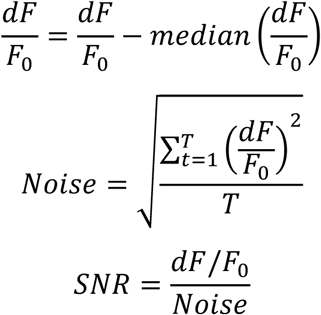

where: *F* is a high-pass filtered fluorescence signal trace, and *F*_0_ is the baseline component of *F*. Both traces were estimated by Volpy, where 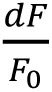 refers to the relative change of *F* regard to *F*_0_. This calculation yielded a time-varying SNR trace, from which the SNR values at spike occurrences were extracted. To assess the long-term stability of GEVI signals, we calculated spike template (average single-spike waveform) at three distinct time intervals (frames 1–300,000, 600,001–900,000, and 1,200,001–1,500,000). Each waveform was obtained by averaging signals from 16 frames (spanning ±8 frames around each spike) at the respective time intervals. In addition to the average waveform, statistical measures such as standard deviation and standard error of the mean (SEM) were computed for individual spike waveforms to assess variability across spikes.

### Whole-cell patch clamp recording in HEK293T

HEK293FT cells were cultured in 24-well plate and transfected with 500 ng of target plasmid DNA using the calcium phosphate protocol as described above. 24 h post-transfection, HEK293FT cells were re-plated on round coverslips coated with Matrigel in DMEM for 1 h at 37°C at a density of 20,000 cells per well in a 24-well plate and incubated for a day at 37°C. Whole-cell patch clamp recording was performed between 36-48 h post-transfection in Tyrode’s solution consisting of (in mM) 125 NaCl, 2 KCl, 3 CaCl_2_, 1 MgCl_2_, 10 HEPES, 30 glucose, pH7.3 (NaOH adjusted) at 320 mOsm; the intracellular solution consisted of (in mM) 135 potassium gluconate, 8 NaCl, 10 HEPES, 4 Mg-ATP, 0.4 Na-GTP, 0.6 MgCl_2_, 0.1 CaCl_2_, pH 7.25 (KOH adjusted) at 295 mOsm.

Data was acquired from HEK293FT cells with access resistance around 5 MΩ, having resting potentials between –50 and –65 mV, membrane resistance > 0.3 GΩ, and holding current (for a holding potential of –70 mV) within ±100 pA. Patch-clamp recordings were acquired via an Axopatch 700B amplifier (Molecular Devices, USA) and Digidata 1440 digitizer (Molecular Devices, USA) in Tyrode’s solution maintained at 23-25°C during experiments using a warmed holding platform (64-1663D, Warner Instruments, USA). Fluorescence imaging was performed on an upright fluorescence microscope (Olympus, Japan), equipped with FITC channel (excitation 475/28 nm from Spectra III LumenCor; emission 594/40 nm) and ORCA-Flash 4.0 V3 sCMOS camera (Hamamatsu), ×5 NA 0.1 and × 40 NA 0.8 objective lenses (Olympus) controlled by NIS Elements AR 5.21.00 (Nikon). The voltage sensitivity was imaged in voltage-clamp mode with a holding potential of--70 mV for 200 ms and then applying voltage steps from –100 mV to + 60 mV for 200 ms at a frame rate of 1 kHz. For quantifying the kinetics, the cells were imaged in voltage-clamp mode with a holding potential of-70 mV for 100 ms and then applying a voltage step of + 30 mV for 200 ms at a frame rate of 2.5 kHz.

### Two-photon scanless voltage imaging in cultured CHO cells

Chinese hamster ovary (CHO) cells (lot#85050302, Sigma-Aldrich, USA) were cultured in Dulbecco’s Modified Eagle’s Medium DMEM-F12 with Glutamax (lot#353170, Thermo-Fischer, USA) supplemented with 10% (v/v) fetal bovine serum and 1 μg/ml penicillin/streptomycin. For experiments, CHO cells were plated onto poly-D-lysine-coated glass coverslips supplemented with 1 μM all*-trans* retinal and transfected using FuGENE® HD (Promega, Madison, USA) the next day. Experiments were carried out 36-45 hours after transfection.

Recordings were performed at 940 nm using a Coherent Chameleon Discovery laser (1.2 W, 80 MHz repetition rate). For scanless two-photon voltage imaging, we projected a temporally-focused holographic spot (∼165 μm^2^, ∼14.5 μm lateral diameter, ∼16 μm axial resolution, 100 mW total power, 0.60 mW/ μm^2^) onto the cell bodies as described before^63,64^. After exiting the laser, the beam was passed through a rotating half-wave plate on a motorized mount, a polarizing beam splitter and a Galilean beam expander giving a size slightly underfilling the spatial light modulator (LCOS X13138-0, 1272 x 1024 pixels, 12.5 μm pitch, Hamamatsu, Japan), which was addressed using the home made software wavefront. The modulated beam was passed through a Fourier lens onto a blazed diffraction grating (600 lines/mm, GR50-0610L11, Thorlabs, USA) and relayed to the focal plane of the objective (CFI APO NI, 40x, NA0.8 numerical aperture, Nikon, Japan). Detection was realized with a simple widefield detection axis using the same objective, a tube lens, and an sCMOS camera (Kinetix, Photometrics, Canada) controlled by micro-manager. The fluorescence passed through a quad-band filter (Chroma ZET405/488/561/640), while the camera was protected from infrared light by a shortpass filter.

Patch pipettes were pulled from thick-walled, fire-polished borosilicate glass capillaries with filament using a P-1000 micropipette puller; resistances ranged between 4-6 MOhm. Signals were amplified (MultiClamp 700B, Molecular Devices, USA), filtered at 10 kHz and digitized at 50 kHz (DigiData1550B, Molecular Devices, USA) using the Clampex software. Whole-cell voltage-clamp recordings were carried out at room temperature; intracellular solution: 110 mM NaCl, 1 mM KCl, 2 mM CaCl_2_, 1 MgCl_2_, 1 mM CsCl and 10 mM HEPES (290 mOsm, pH 7.2); extracellular solution: 140 mM NaCl, 2.5 mM KCl, 2 mM CaCl_2_, 1 mM MgCl_2_, 12.5 mM D-Glucose, 10 mM HEPES (310 mOsm, pH 7.2). The liquid junction potential was calculated to be +1.2 mV and not corrected. Pipette capacitance was compensated in cell-attached configuration before break-in. Series resistance was not compensated. R_m_ was usually in the GOhm range (never below 600 MOhm) while R_a_ was below 15 MOhm. During the experiment cells were patched and after obtaining whole-cell configuration, the membrane potential was stepped from – 60 mV to + 40 mV (voltage clamp mode) for 100 ms, 10 ms and five times 2 ms, while fluorescent changes were imaged simultaneously with the sCMOS camera under holographic two-photon illumination at a frame rate of 1006 Hz.

Data was acquired in TIFF format and further analyzed using Fiji and custom Python scripts. To obtain a binary mask, a maximum intensity projection of the stack was created, and cells were semi-manually segmented, supported by Otsu-thresholding based on intensity; intense regions (for a few cells) located in the center of the cells were ascribed to imperfect membrane targeting and excluded from the masks. The raw fluorescence trace was calculated as the spatial average fluorescence over the mask and background corrected by subtraction the background trace, obtained as the spatial average fluorescence over 50 not illuminated columns of the frame. The background-corrected trace (excluding the frames were the voltage was stepped) was then used to calculate a rolling F_0_ by applying a gaussian filter (sigma=200), which was the basis for calculating ΔF/F = (F-F_0_)/F_0_ and SNR = (F-F_0_)/SD_F(-60 mV)_ traces used for further analysis. All values for ΔF/F and SNR were then extracted from the respective traces and time points. For the 100-ms step, the values were averaged over the last 50 ms of the step, while for the 2-ms steps, the maximum values were extracted.

### Animals

All animal maintenance and experimental procedures were conducted according to institutional and ethical guidelines for animal welfare, and all animal studies were approved by the Institutional Animal Care and Use Committee (IACUC) of Westlake University, Hangzhou (animal protocol #KP-19-044) or the Hebrew University of Jerusalem corresponding to the location of the performed experiments. For Supplementary Fig. 7a, C57BL/6J mice aged 8-18 weeks were used. For Figure 3, **Supplementary Figure 1, 2, and Supplementary Figure 7b**, VIP-Cre (JAX# 010908) and C57BL/6J mice were used, regardless of sex.

### One-photon voltage imaging in awake mice in the primary visual cortex

Stereotactic injection and cranial window surgery were conducted as previously described^65^. Briefly, VIP-Cre mice (JAX#010908, USA) aged 8 weeks were anesthetized using 4% isoflurane (induction) and maintained using 1.5-2.0% on a 37°C heating pad for the entire surgery procedure. After head-fixing the mouse on a stereotaxic apparatus (RWD Life Science, USA), hair on the scalp was removed using depilatory cream, and eyes were covered with erythromycin ophthalmic eye ointment. Then, a craniotomy of ∼3 mm in diameter was created over the visual cortex (AP: −3.4 mm, ML: 2.1 mm). 100 nl of either AAV2/9-CAG-DIO-ElectraOFF-Kv2.1 or AAV2/9-CAG-DIO-Ace-mNeon2-Kv2.1 (>10^13^ vg/mL, Shanghai Sunbio Medical Biotechnology, China) was injected at 50nl/min into the primary visual cortex (AP:-2.5 mm, ML: 2.5 mm), 200-μm depth. A total of 5-6 injections were made at the left visual cortex (ΔAP: 0 to-1mm, ΔML: ±1 mm, centering at AP:-2.5 mm, ML: 2.5 mm). For the imaging of pyramidal neurons, wild-type C57BL/6J mice were used, and a mixture of AAV2/9-CAG-DIO-ElectraOFF-Kv2.1 (final titer 10^13^ vg/mL) and AAV2/9-CaMKIIa-Cre (final titer 10^10^ vg/mL) was injected. The stereotactic injection was performed the same as for VIP-Cre. Right after stereotactic virus injection, a circular coverslip (3 mm in diameter, 0.1 mm in thickness, Luoyang GULUO Glass, USA) was attached to the dura surface and Kwik-sil adhesive (World Precision Instruments, USA) was applied around the edges of the imaging window to anchor the imaging window to the skull. Then, dental cement (C&B Metabond, Parkell, USA) was applied to cover all skull surface. A custom titanium headplate was then placed above the cement. After the dental cement solidified, a layer of denture base resin was applied on top of the dental cement to consolidate the entire window and left for 5-10 minutes to dry. The mouse was allowed to recover for at least 3-4 days after surgery.

### Wide-field one-photon imaging

After 3-4 weeks post-injection and window implantation, the mice were ready for imaging. Prior to image acquisition, mice were anesthetized using 2% isoflurane and head-fixed onto a custom stage. Wide-field imaging was then performed by using an epifluorescence microscope (BX51WI,Olympus, Japan), a 20x water-immersion objective (NA0.95, morphology imaging) and a sCMOS camera (OrcaFlash4.0v2, Hamamatsu). Epi-illumination was provided by a 473 nm laser (Changchun New Industries Optoelectronics Tech, China) adjusted at 50 mW/mm^2^ and 100 mW/mm^2^, respectively; emission light was filtered through a multiband bandpass filter (ZT405/473/561rpc, Chroma, USA). Typically, neurons from 130-200 μm below the pia were recorded. To ensure efficient image transfer and prolonged recording, we conducted consecutive imaging trials. Briefly, 19140 (x*y, 80-180 × 80 pixels, equivalent to 104-234×104 μm, 4×4 binning) frames were acquired at 638 Hz (30s continuous recording), followed by a timed 30s-period without imaging or light exposure. The recording remained consecutive until one of the following situations took place: for each ten trials conducted, we monitor whether neurons became non-spiking over at least five sessions and/or the fluorescence baseline of neurons was blended with background. Those inter-trial checks could contribute to a 30-sec to 1-min delay before the next set of ten-trial recording; however, they initiated a limited effect on the overall tendency of fluorescence baseline drop and thus drop in spike SNR level in the long term. Over the course, recordings were conducted within 4 to 7 weeks post-viral injection.

### One-photon voltage imaging in awake mice in CA1 of hippocampus

Virus injections and surgical procedures followed protocols similar to those described previously^66^. Briefly, mice at age of 8-18 weeks were anesthetized with 2% isoflurane for induction and maintained at 1.0-1.5% isoflurane during surgery. The surgical site was exposed by retracting the scalp, and the skull surface was rinsed and dried. A 3-mm diameter craniotomy was performed over the left dorsal CA1 region (ML: 1.8 mm, AP: 2.3 mm) using a 3-mm biopsy punch (Miltex). The dura mater was carefully removed, and the cortex was aspirated under continuous saline irrigation. The external capsule was then exposed, and a small central portion was delicately removed to expose the CA1 area.

Viral infection was performed by generating viral eluting cover glasses as described before for GRIN lenses^67^. In brief, 1 μl of AAV9:CAG-DiO-ElectraOn virus (final titer 1 × 10^12 GC/ml) plus 0.8 μl of rAAV9-hSyn-Cre (final titer 1.6 x 10^10 GC/ml) was mixed with 2 μl of 0.6% CMC 1.4% Trehalose solution (w/v) and was placed on the cover glass of a custom-made cannula. The cannula consists of a 2.0-mm segment of 3-mm diameter stainless steel tube (MicroGroup, USA) capped with a no.1 round cover glass using UV curable glue (lot#NOA81, Norland, USA). The cannula was left to air-dry for 30 minutes and then implanted and secured to the skull with 3M Vetbond tissue adhesive. After the cannula was secured, a custom-made titanium head plate was glued around the cannula, and any exposed skull was covered with dental cement (C&B Superbond, Sun Medical, USA). Animals were returned to the cage for recovery and treated with meloxicam (5 mg kg-1) for 2 days.

In vivo voltage imaging was performed 2-8 weeks following surgery. Mice were anesthetized with 2% isoflurane for induction and maintained at approximately 1% isoflurane during imaging. Imaging was conducted on a widefield epifluorescence module (Thorlabs, USA). The microscope was also equipped with a custom holography module designed and built by Thorlabs Imaging Systems Inc. according to our specifications. The module included a 120 mW CW 488 nm laser (Coherent, USA) that was fiber-coupled and patterned using a spatial light modulator (SLM, EXULUS-HD2, Thorlabs, USA). The beam was expanded, collimated, passed through a ¼ wave plate, targeted to half of the SLM, and relayed to the objective back aperture. The zero’s order was blocked in an intermediate image plane using a glass with a deposited gold point. Mice were imaged using a 16X 0.8 NA 3-mm working distance objective (Nikon, N16XLWD-PF) and a 252 mm tube lens (20.1X magnification). The laser was attenuated using a MEMS-based electronic attenuator (V450A, Thorlabs), and cells were imaged at 160 mW/mm^2^. Images were collected on a scientific CMOS camera (ORCA-Fusion, Hamamatsu, Japan) at a sampling rate of 500Hz.

### Data extraction and analysis for *in vivo* one-photon images

For batch processing of *in vivo* one-photon image data and the extraction of neuronal signals across multiple trials, we adapted and modified previously described methods^68,69^. The introduced modifications are described below.

*Motion correction:* we modified the algorithm to load image files in.nd2 format, and a rigid registration algorithm based on Fast Fourier transformation was used for motion correction. The necessary frame-to-frame shifts were computed via cross-correlation, and the corrected images were obtained using bilinear interpolation to remove motion artifacts.

*Background correction and ROI trace extraction:* to remove global background fluctuations and bleaching, an 8×8 spatial binning was first applied to the raw image. Singular value decomposition (SVD) was then used to extract principal components, and the first principal component was subtracted from the raw images by linear regression. ROIs corresponding to individual neurons were then manually selected and had raw traces extracted using the mean value. The local background signal was then estimated from an annulus extending 15 pixels around each ROI and was subtracted from the trace extracted to remove background fluorescence.

*Trace denoising and spike detection:* The background-corrected fluorescence traces were high-pass filtered with a third-order Butterworth filter at 10 Hz to remove low-frequency baseline fluctuations. The resulting signals were centered by subtracting their median values. Then, initial estimates of the spike temporal position were identified by applying a threshold with standard deviation (SD) > 3.5 from the baseline noise level. For ElectraOFF and Ace-mNeon2, the positive part of the trace was treated as noise, whereas negative signals were treated as noise for ElectraON. A spike template waveform was generated by aligning and averaging these putative events within a 10-ms window, centered at spike position. A whitened matched filter^70^ was then applied to the fluorescence trace to emphasize features by temporal matching, followed by an adaptive threshold (minimal SD = 3.5) to refine spike detection in the matched-filtered trace. Fluorescence traces displayed in figures, as well as those used for signal-to-noise ratio (SNR) calculations, were not processed with the high-pass filter or whitened matched filter.

Since Gaussian shot noise can randomly introduce fluctuations in both positive and negative directions, spike detection was enabled only if the total number of detected spikes exceeded the number of events in the noise (opposite) direction. For trial-based recording, we observed that SNR decreased significantly for Ace-mNeon2 and spike number per session also decreased through time. To maximize the chance of assessing SNR from spikes (but not noise) in longer recording sessions for both probes, we extracted and used peak SNR from individual image sessions to report spike SNR through time in vivo. Based on experimental observation, a threshold of SD > 3.5 could potentially introduce 2–4 false-positive spikes per trial of recording, aligning with the predicted number of spikes caused by noise (PSN) from Gaussian noise (when SD = 3.5, PSN = 4:

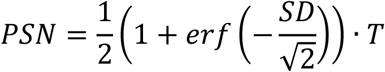

where *erf* denotes the Gauss error function, SD is the number of standard deviations from the baseline noise level chosen as the threshold for spike detection, and *T* is the number of time frames. We thereby further manually inspected all traces to make sure that any imaging trial with a dominance of false-positive spikes was excluded from the peak SNR analysis. Meanwhile, if no spikes surpassed the initial threshold for spike template generation or the total number of detected spikes is smaller than the number of events in the noise (opposite) direction, the ROI was categorized as non-spiking and was not analyzed further for that session.

## Data availability

The most essential raw datasets, including source files for supplementary figures and raw unprocessed images, are available at figshare (https://figshare.com/articles/media/Representative_raw_data_files_for_4_voltage_sensors_pAce_Ace-mNeon2_ElectraON_and_ElectraOFF/28935143). The remaining files are available from the corresponding authors upon request. All plasmids used in this study are available from WeKwikGene (https://wekwikgene.wllsb.edu.cn/) and Addgene. Source data are provided with this paper.

## Code availability

Custom codes used in this study for voltage imaging data analysis are available at https://github.com/zhanghanbin598/Voltage-imaging-codes-for-Electras and https://github.com/Shihao-Neuro/GEVI.

## Acknowledgements

We thank Stavrini Papadaki and Xun Gao from Westlake Laboratory for verifying all plasmid sequences and depositing them to WeKwikGene. We thank the Laboratory Animal Resources Center and Biomedical Research Core Facilities at Westlake University for their help with technical support. We thank Lu Bai from Kai Wang lab for her help with performing the Matlabcodes to analyze the voltage imaging data. We thank Hui Xiong from Bo Li lab for his suggestions on whole-cell patch clamp. This work was supported by start-up funding from the Foundation of Westlake University, Westlake Laboratory of Life Sciences and Biomedicine, National Natural Science Foundation of China grant 32171093, ‘Pioneer’ and ‘Leading Goose’ R&D Program of Zhejiang 2024SSYS0031 to K.D.P, and National Natural Science Foundation of China grant 323B200334 (H. Z.).

## Contributions

K.D.P. initiated the project and, together with H.Z. and S.Z., made high-level designs and plans and interpreted the data. H.Z. and S.Z. developed the ElectraON and ElectraOFF indicators and characterized them in mammalian cell culture. T.P.K., O.M.S., and F.V.S. developed mBaoJin(3M). H.Z. performed whole-cell patch clamp experiments. S.Z. performed one-photon *in vivo* imaging inmouse primary visual cortex. S. B.-S., and Y.A. performed *in vivo* imaging in mouse hippocampus CA1. H.Z., S.Z., and C.G. prepared figures. H.Z., S.Z., and M.E. performed data processing and analysis. C.G. and V.E. characterized indicators under 2P excitation. H.Z., S.Z., and K.D.P. wrote the paper with contributions from all of the authors. K.D.P. oversaw all aspects of the project.

## Supplementary Information

**Supplementary Table 1.** List of the reported long-term voltage imaging experiments.

| Sensor | Cell types/<br>Organism | Imaging settings | Application | Duration | Refs |
| --- | --- | --- | --- | --- | --- |
| pAce | L1, L2/3 neurons in primary visual cortex and CA1 of mice<br>CaMKII, VIP positive neurons in mice<br>MBON-γ1pedc>$\alpha/\beta$ and PPL1-g2a'1 neurons in Drosophila flies | Wide-field one-photon, 400-600 Hz in mouse<br>1kHz in fly;<br>The illumination intensity was ~25 mW/mm<sup>2</sup> in V1 and ~25 to 75 mW/mm<sup>2</sup> in CA1.<br>The 30-min continuous imaging session in Drosophila use ~5 mW/mm<sup>2</sup> laser power. | 1) Representative recordings from PPL1-g2a'1 of flies showing odor-evoked spiking elicited by 5-s delivery of 5% isoamyl acetate<br>2) Concurrent 0.6 kHz DUPLEX imaging and LFP recordings in hippocampal CA1<br>3) Dual population recordings in V1 and CA1 during state transitions.<br>4) Simultaneous dual-polarity and dual-color imaging capture the voltage dynamics of three distinct targeted subtypes or cell subclasses in awake mice. | From **several seconds of imaging to several minutes** of imaging.<br>The longest imaging is a **30-minute** continuous imaging session in Drosophila flies. | <sup>72</sup> |
| Ace-mNeonGree n2 | L1, L2/3 neurons in primary visual cortex and CA1 of mice<br>NDNF, VIP, SST, and CaMKII-positive neurons in mice<br>PPL1-g2a'1 neurons of flies | Widefield one-photon,<br>400-600Hz in mice<br>The illumination intensity at the sample plane for all indicators was ~25 mW/mm<sup>2</sup> in V1 and ~25 to 75 mW/mm<sup>2</sup> in CA1. | 1) Representative recordings from PPL1-g2a'1 of flies showing odor-evoked spiking elicited by 5-s delivery of 5% isoamyl acetate (IAA).<br>2) Example visual responses from layer 2/3 PNs to drifting gratings presented at eight different orientations, acquired using Ace-mNeon2 in an awake mouse.<br>3) Ace-mNeon2 voltage imaging of different neuronal subtypes in mice during air puff delivery.<br>4) Concurrent 0.6 kHz DUPLEX imaging and LFP recordings in hippocampal CA1<br>5) Dual population recordings in V1 and CA1 during state transitions.<br>6) Simultaneous dual-polarity and dual-color imaging capture the voltage dynamics of three distinct targeted subtypes or cell subclasses in awake mice. | From **several seconds of imaging to several minutes** of imaging.<br>The longest imaging is 5 min-long in vivo spike imaging using Ace-mNeon2 in an awake mouse | <sup>72</sup> |
| JEDI-2P | mouse retina<br>L2/3 neurons in primary visual cortex of mice | Two-photon<br>Imaging was conducted at 440 Hz | 1) JEDI-2P captures voltage responses to changes in visual stimuli frequency and contrast in isolated mouse retina.<br>2) Imaged axonal projections of L2 cells, non-spiking neuron postsynaptic to photoreceptors (R1–R6) in Drosophila.<br>3) Voltage dynamics in layer 2/3 cells of the visual cortex of mice.<br>4) more than >40 min of continuous ULoVE optical recording<br>5) Baseline-corrected fluorescence signals from two neurons of layer 2/3 recorded simultaneously for 15.4 min. | Retaining 72.4% ± 0.1% of its fluorescence over **30 min** of imaging with 34 mW of power<br>More than **>40 min** of continuous | <sup>73</sup> |
|  |  |  |  | ULoVE optical recording |  |
| JEDI-1P | L2/3 neurons in primary visual cortex of mice | Widefield one-photon:<br>~36 mW/mm<sup>2</sup><br>0.05 mW/mm<sup>2</sup> | 1) Pan-cortical population JEDI-1P-Kv signals correlate with local field potentials (**no single cell resolution**).<br>2) Population imaging of JEDI-1P-Kv reports rapid voltage responses to high-frequency stimuli (**no single cell resolution**). | Under 0.05 mW/mm<sup>2</sup>, more than 5 h of imaging over 19 days—10 imaging sessions of one hundred 20-s trials—JEDI-1P-Kv conserved $77.4 \pm 1.3\%$ of its fluorescence.<br>Under continuous illumination, JEDI-1P-Kv's intensity decreased to $95.8 \pm 4.2\%$ after 1 h. | <sup>74</sup> |
| ASAP4b and ASAP4e | hippocampal SST+ interneurons<br>hippocampal place cells<br>cortical pyramidal neurons | Two-photon<br>1,000 fps,<br>35mW/mm<sup>2</sup>, 50<br>mW/mm<sup>2</sup>,<br>100mW/mm<sup>2</sup>, 150<br>mW/mm<sup>2</sup><br>One photo imaging<br>1,000 fps,<br>100 mW/mm<sup>2</sup> | Simultaneous voltage and calcium imaging in cortical pyramidal neurons in vivo. | With minimal modifications to a commercial resonant-scanning two-photon microscope, we could continuously image rectangular fields of view containing >50 cells at 989 fps for **tens of minutes** | <sup>75</sup> |
| ASAP5 | Mice<br>fruit flies | One-photon imaging<br>500 fps, ~46<br>mW/mm<sup>2</sup> | 1)Single-trial imaging of graded and action potentials in fruit flies<br>2) Single-trial two-photon imaging of supra- and subthreshold activities in awake mice<br>3)Voltage imaging of dendrosomatic propagation of mEPSPs | **2-min** 500 fps GEVI recordings | <sup>76</sup> |
| Voltron2 | Zebrafish | For flies, Images were acquired at 800 Hz, 2-11 mW/mm<sup>2</sup>. | 1) In vivo voltage imaging of olfactory sensory neurons in zebrafish;<br>2) Imaging of voltage activity in vivo in flies; | For Zebrafish, fluorescent signal was | <sup>77</sup> |
|  | Flies<br>mouse hippocampus, visual cortex, and motor cortex | In mice, the illumination intensity was ~70–140 mW/mm<sup>2</sup> for Parvalbumin (PV) interneurons in mouse hippocampus; 18.5 mW/mm<sup>2</sup> in mouse visual cortex; <20 mW/mm<sup>2</sup> mouse anterior lateral motor cortex | 3) In vivo voltage imaging in mouse hippocampus, visual cortex, and motor cortex | recorded for **5 minutes**<br>Imaging Parvalbumin (PV) interneurons in mouse hippocampus for **3 minutes**.<br>At the aforementioned illumination levels, spiking activity was detectable for over 20 minutes in PPL1-γ1pedc and over **10 minutes** in MBON-γ1pedc>$\alpha/\beta$. |  |
| SpikeyGi2 | Mouse, L2/3 neurons in the primary somatosensory cortex (S1) of mice, the anterior olfactory nucleus, and main olfactory bulb neurons | Two-photon, 1 kHz frame rate, 400x400 μm FOV), low-photon flux (~30 mW per beamlet, 8 beamlets), 920 nm. | Simultaneous high-speed deep-tissue imaging of >100 neurons, spike detection below shot-noise limit using DeepVID denoising, tested with whisker stimulation. | Sustained imaging over 1 hour in awake behaving mice with minimal photobleaching under intermittent imaging, no significant photodamage detected. | 24 |

---

**Supplementary Figure 1.**
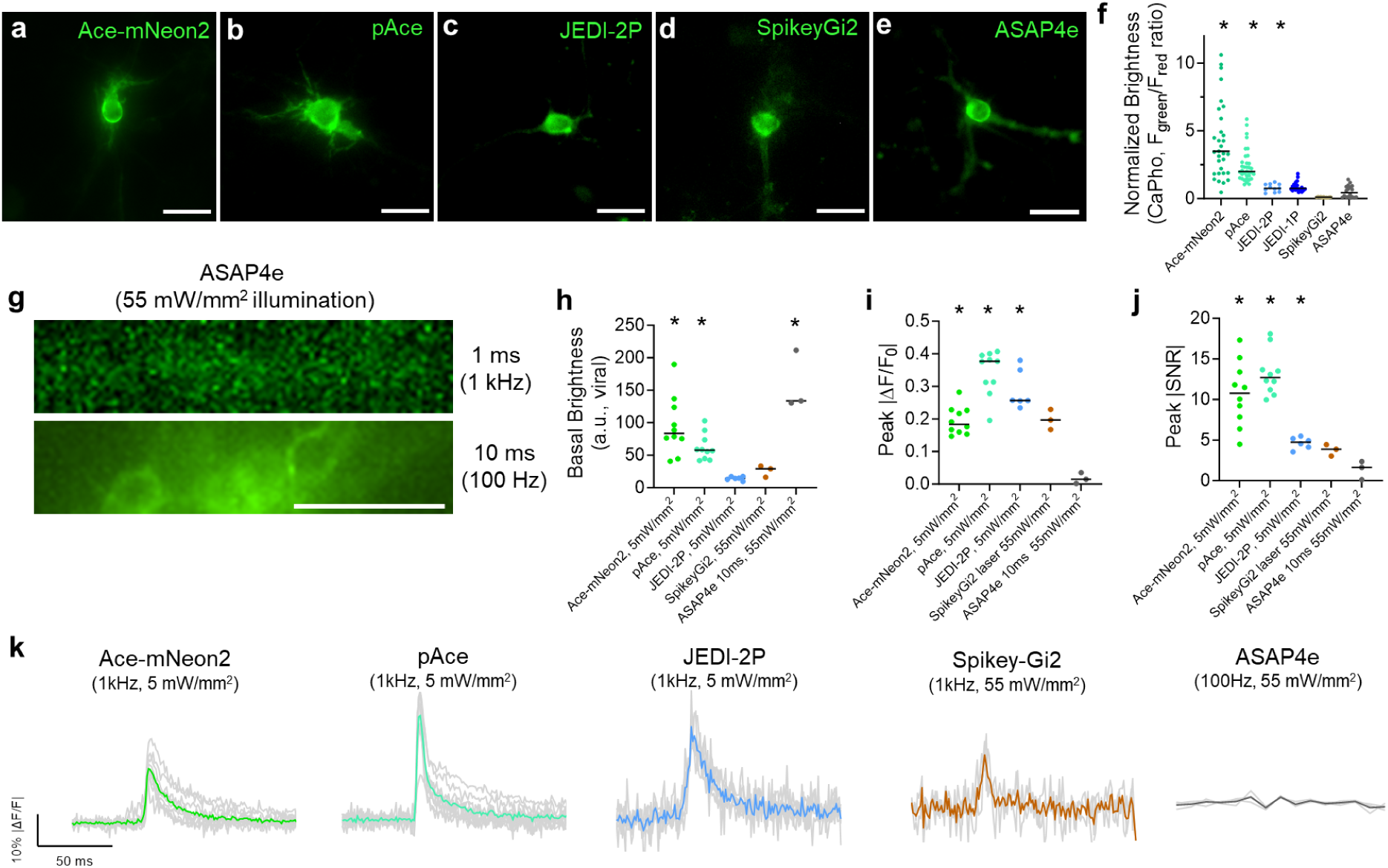
Quantitative assessment of genetically encoded voltage indicators in primary hippocampal mouse neuronal culture. **a-e.** Representative wide-field images of transfected neurons expressing soma-targeted (a) Ace-mNeon2, (b) pAce, (c) JEDI-2P, (d) SpikeyGi2, and (e) ASAP4e. **f.** Comparison of brightness among GEVIs in calcium phosphate-transfected neuronal culture. n = 33 neurons for Ace-mNeon2, n = 35 neurons for pAce, n = 11 neurons for JEDI-2P, n = 11 neurons for SpikeyGi2, n= 18 neurons for Ace-mNeon2, from 2 cultures. The green fluorescence was normalized by fluorescence from membrane-localized fusion red, reported as green-to-red fluorescence ratio. **g.** Maximum projection image showing viral transduced neurons expressing ASAP4e, sampled under 100Hz and 1kHz. Under 1 kHz, the baseline fluorescence of ASAP4e could not be resolved. **h.** Basal brightness comparison between GEVIs in viral transduced neuronal culture. **i-j.** Absolute peak ΔF/F0 (i) and SNR (j) comparison between GEVIs in viral transduced neuronal culture. For SpikeyGi2 and ASAP4e, voltage recordings were performed at 55mW/mm^2^, and a 100 Hz sampling rate was applied to ASAP4e to detect fluorescence baseline. **k.** Averaged spike waveform generated in response to 1-AP electrical stimulation GEVIs in transduced neuronal culture. Scale bar, 50μm.

**Supplementary Figure 2.**
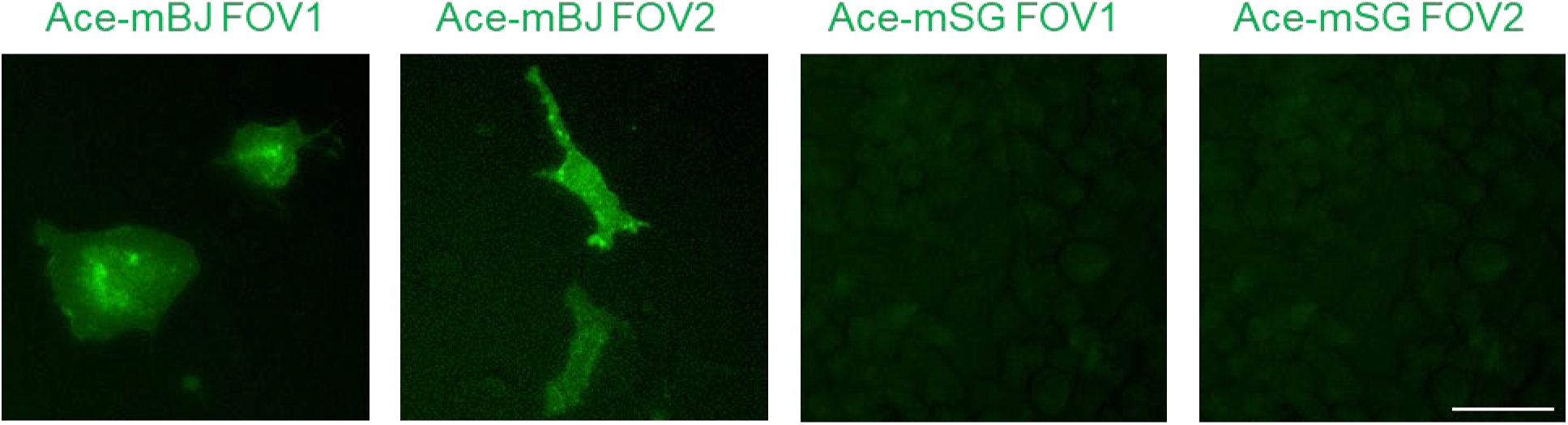
Expression of Ace-mBJ and Ace-mSG in HeLa cells. Wide-field images of HeLa cells expressing Ace-mBJ and Ace-mSG under the same laser power; expression time 48 h.

**Supplementary Figure 3.**
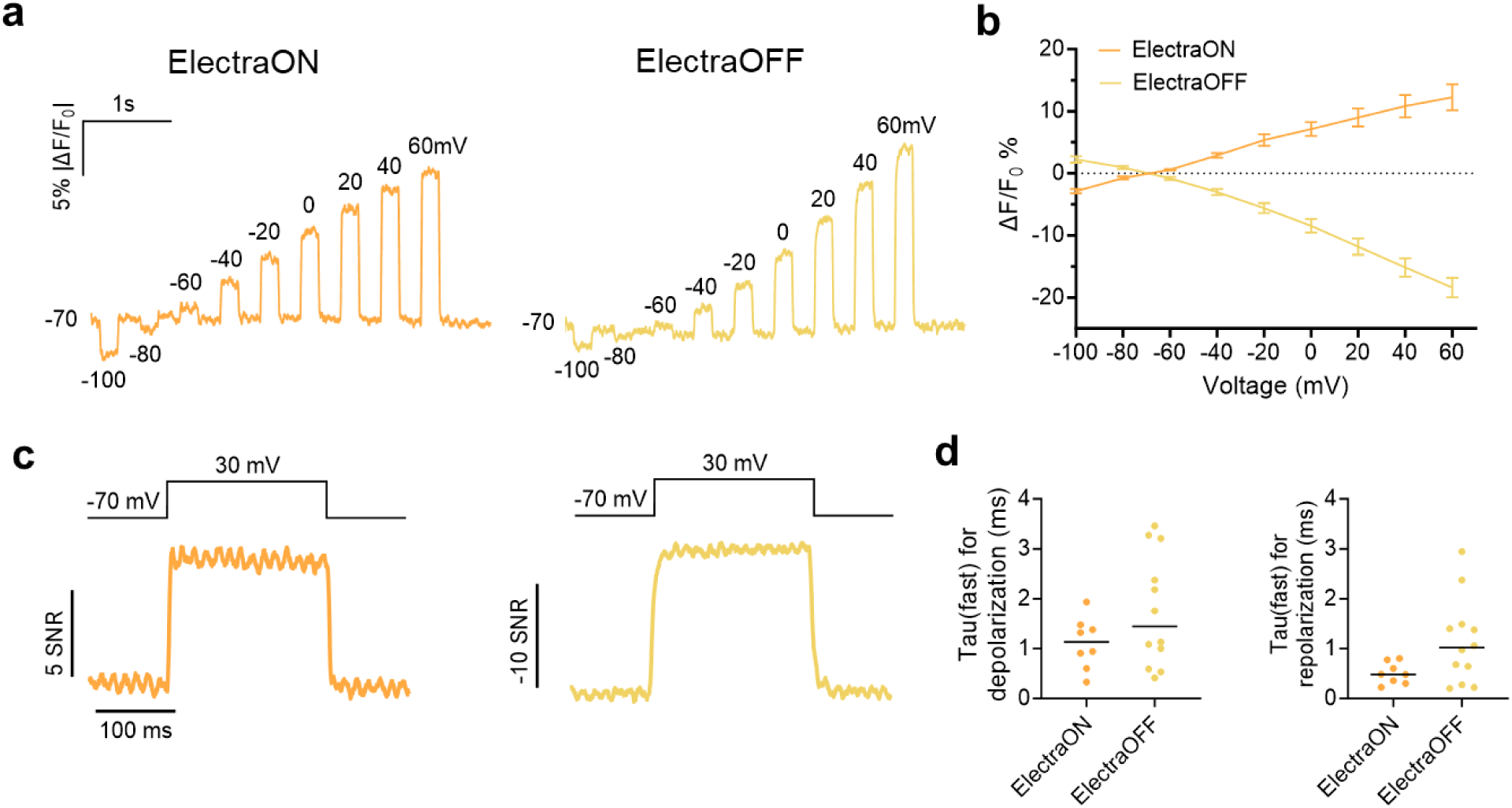
Electrophysiological characterization of ElectraON and ElectraOFF in HEK293T cells using whole-cell patch clamp. **a.** Steady-state responses to voltage commands in HEK cells of ElectraON and ElectraOFF under 20 mW/mm² laser power, normalized to −70 mV (n = 6 cells for ElectraON, n = 5 cells for ElectraOFF from three cultures). Image acquisition rate: 1 kHz. **b.** F-V curves of ElectraON and ElectraOFF. n = 6 cells for ElectraON, n = 5 cells for ElectraOFF. Error bars represent s.e.m. **c.** Representative fluorescence response of ElectraON and ElectraOFF in HEK cells to a 100-mV change in voltage clamp. Image acquisition rate: 2.5 kHz. **d.** Kinetics (τfast) of ElectraON and ElectraOFF were characterized in the depolarization (-70 mV to +30 mV) and repolarization (+30 mV to-70 mV) stages.

**Supplementary Figure 4.**
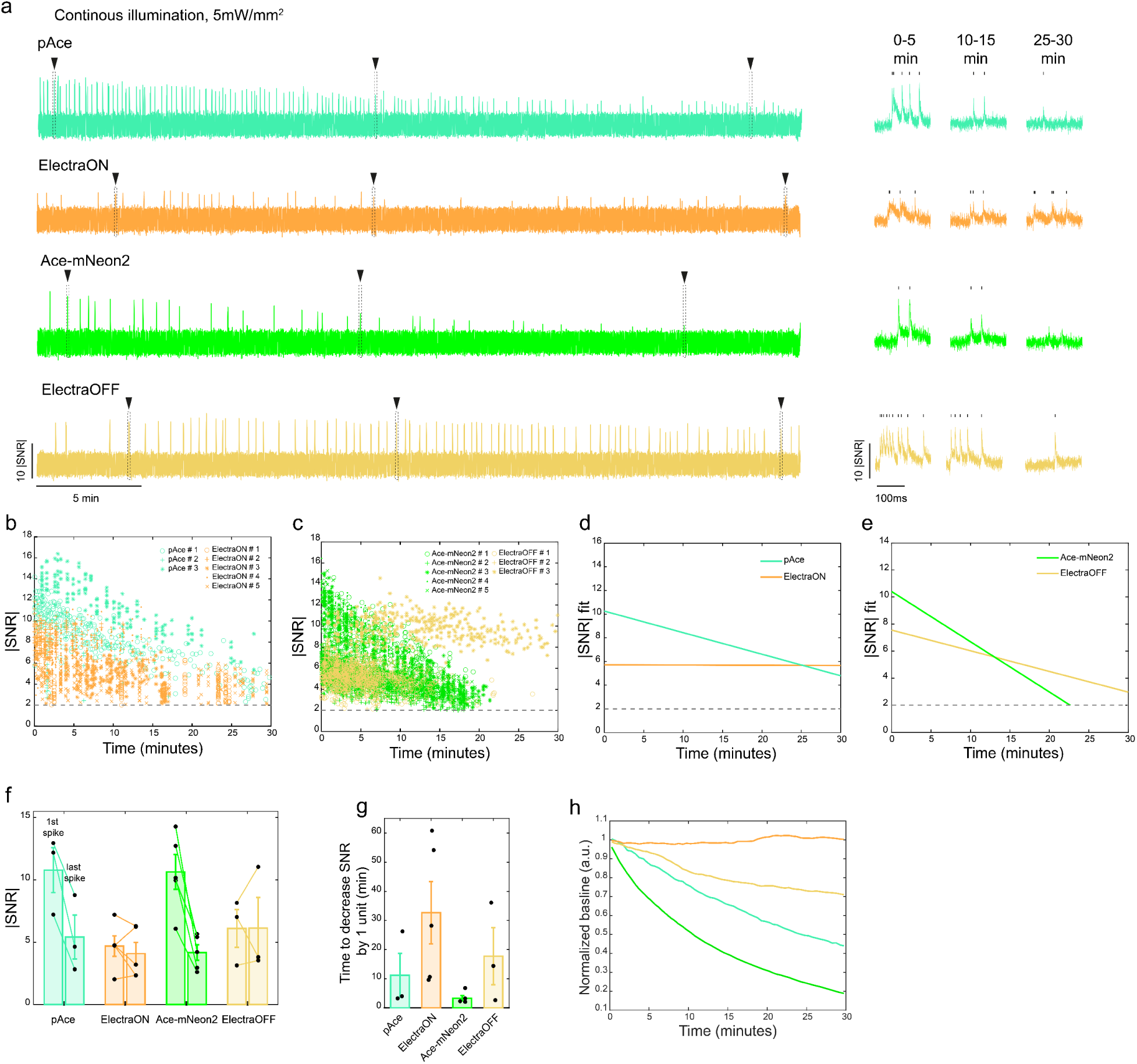
30-minute continuous recording of GEVIs in cultured primary hippocampal neurons. **a.** Example full 30-min raw fluorescence trace of absolute SNR and zoom-in-view of 200ms-trace snippets for GEVIs sampled from the first 5 min, middle 5 min and last 5 min of recording. **b.** Individual SNR values of spikes identified for pAce-and ElectraON-expressing neurons from 30-minute continous recording. **c.** Same as to b, but for Ace-mNeon2 and ElectraOFF. **d.** Fitted average of the SNR values for pAce-and ElectraON-expressing neurons from 30-minute continous recording. **e.** Same as to d, but for Ace-mNeon2 and ElectraOFF. **f.** Absolute SNR value of the first and last spike identified for GEVIs. **g.** Time required for SNR to drop by 1 unit for GEVIs from a 30-minute continuous recording. **h.** Curve showing the photobleaching of GEVI fluorescence baseline from a 30-minute continuous recording. n = 3 neurons for pAce, 5 neurons for ElectraON, 5 neurons for Ace-mNeon2 and 3 neurons for ElectraOFF, from 3 independent cultures.

**Supplementary Figure 5.**
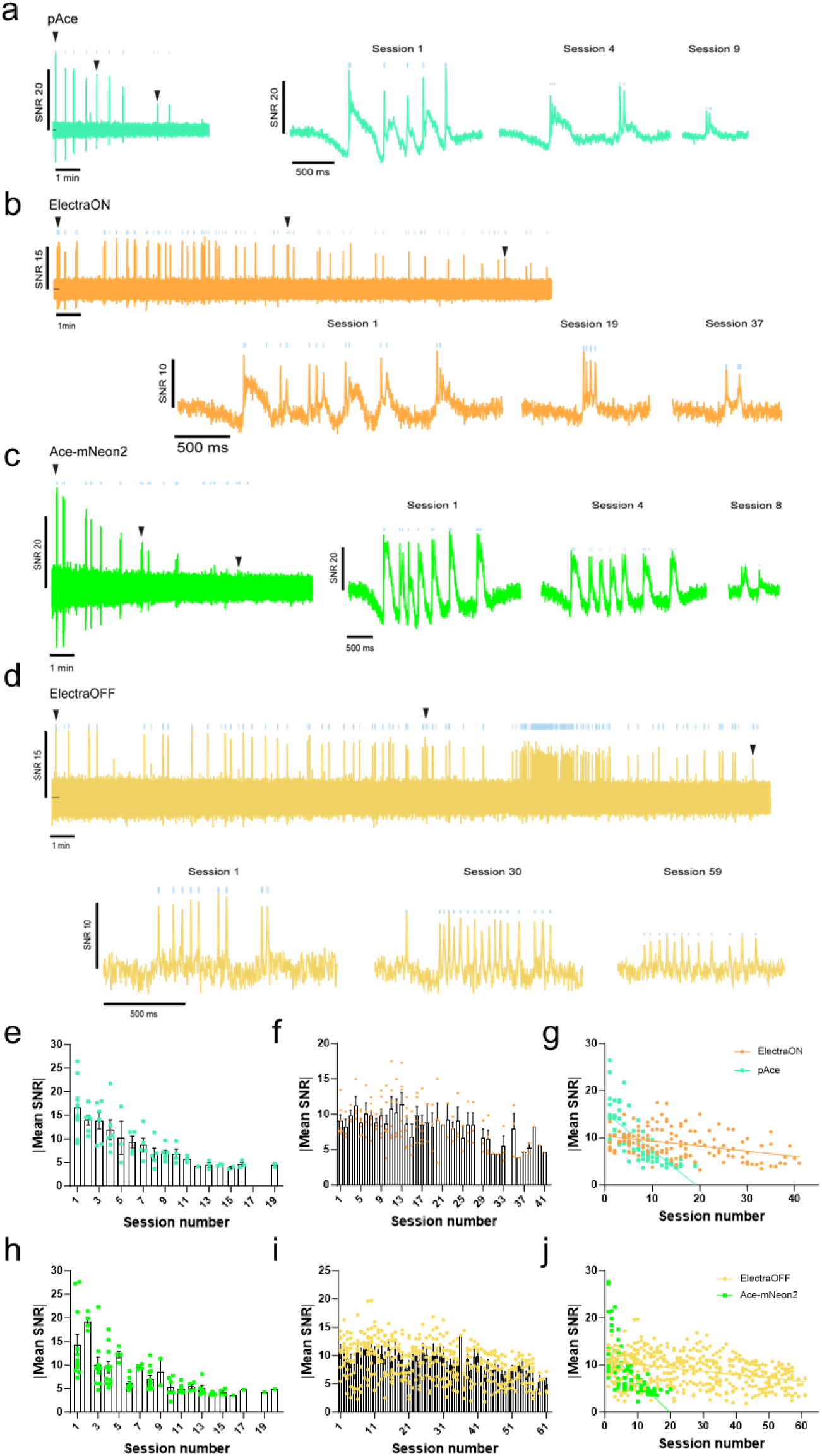
Long-term voltage imaging of cultured neurons expressing (p)Ace-based voltage sensors at 25mW/mm^2^. **a.** Left panel: overview of all end-to-end 30-second traces of neuronal activity recorded from pAce-expressing neurons. Right panels: Three zoomed-in traces corresponding to the arrow-indicated regions (session1, session5 and session 8) in the overview trace, demonstrating local activity patterns and SNR level. **b-d.** Same as a but for ElectraON(b), Ace-mNeon2(c) and ElectraOFF(d). For ElectraON, zoomed-in traces corresponding to session1, session12 and session 22 were shown; For Ace-mNeon2, zoomed-in traces corresponding to session1, session3 and session 6 were shown; For ElectraOFF, zoomed-in session1, session28 and session 56 were shown. **e-f.** Absolute mean SNR value of spikes per neuron calculated for pAce(e) and ElectraON(f) across imaging sessions. n=7 neurons for pAce and n = 13 neurons for ElectraON. **g.** Linear fitting of absolute mean SNR values for pAce and ElectraON in e-f. **h-i.** Same as e-f but for Ace-mNeon2(h) and ElectraOFF(i). n=10 neurons for Ace-mNeon2 and n = 14 neurons for ElectraOFF. **j.** Linear fitting of absolute mean SNR values for Ace-mNeon2 and ElectraOFF in h-i.

**Supplementary Figure 6.**
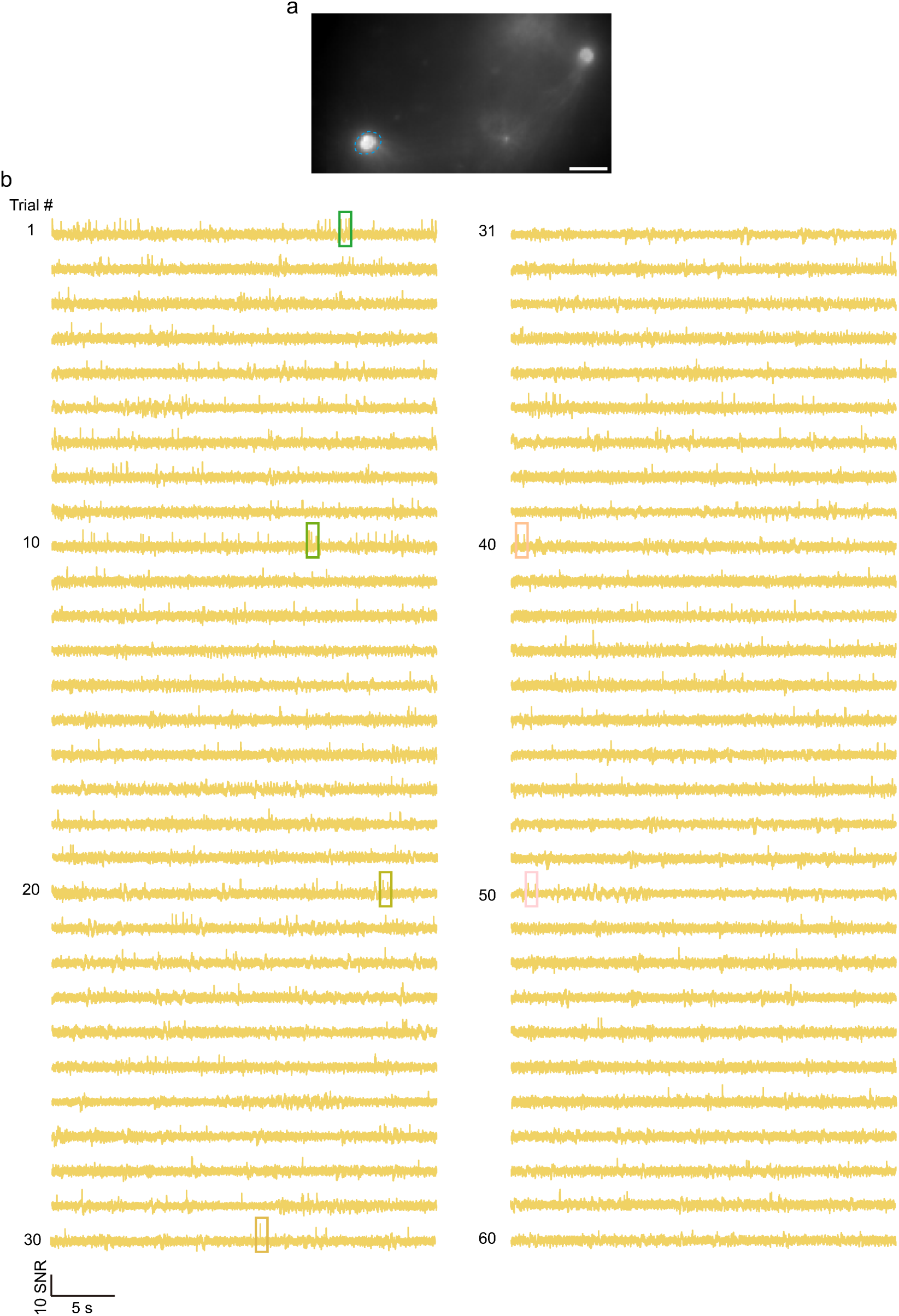
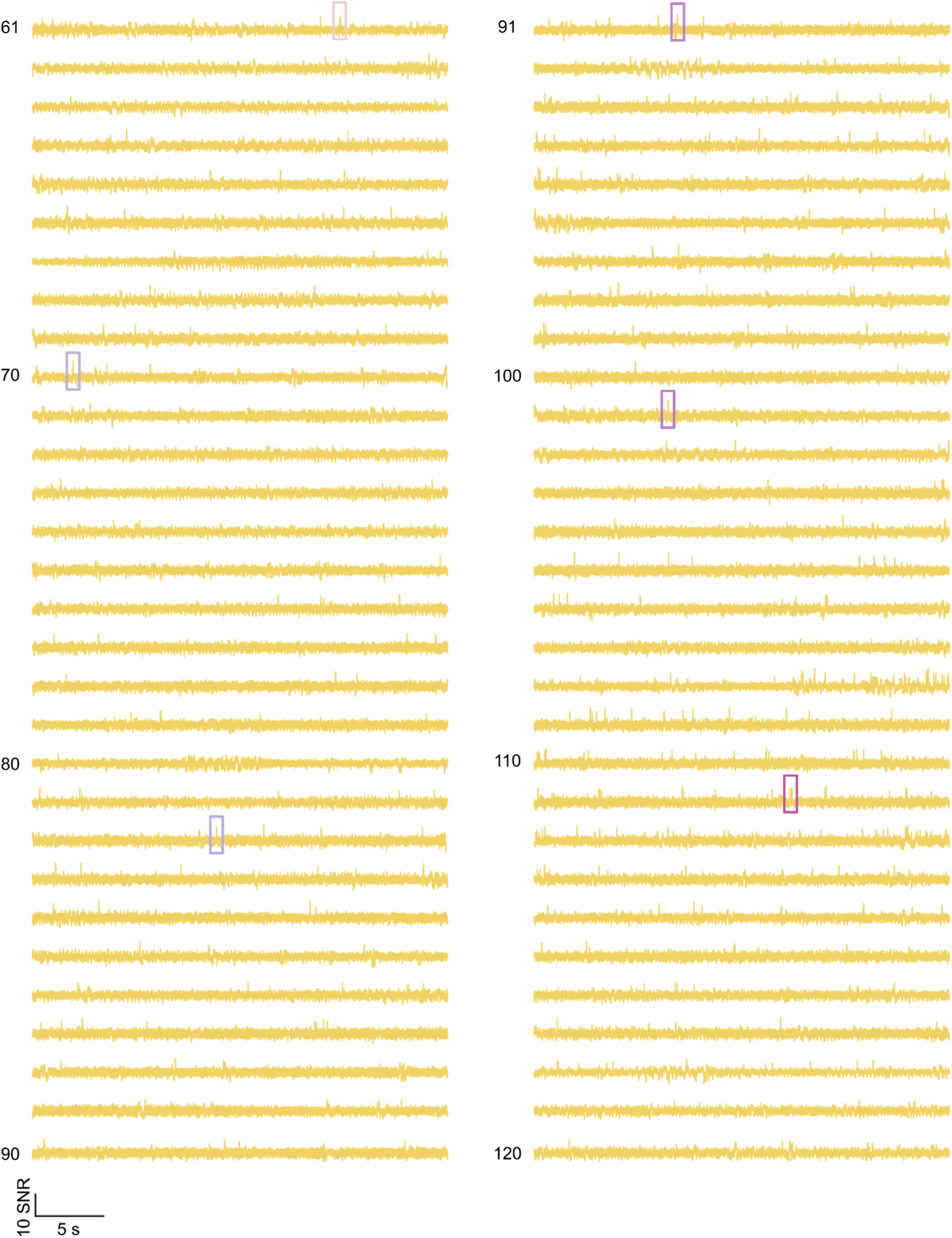
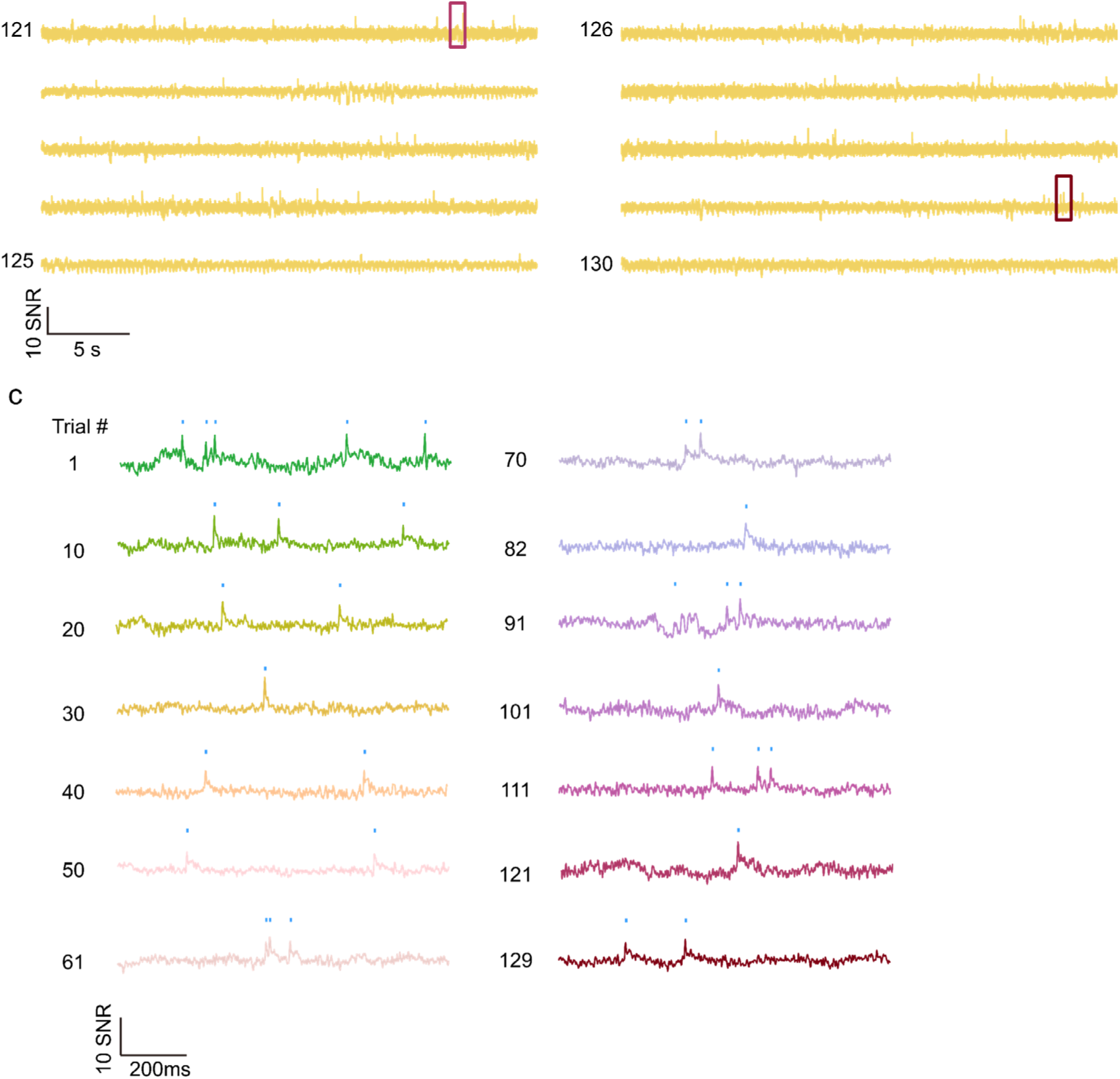
Hour-long full voltage recording of ElectraOFF expressing VIP+ interneurons in V1 using one-photon microscopy. **a.** Average projection image of ElectraOFF-expressing VIP+ interneurons in V1 with the target neuronal ROI highlighted in blue. **b.** 30-sec raw fluorescence trace of the neuron shown in a for a total of 65-min recording (equivalent to 130 imaging trials). **c.** 1-sec trace snippets sampled from every 4-5 min of recording from the colored box in b. Scale bar = 25μm.

**Supplementary Figure 7.**
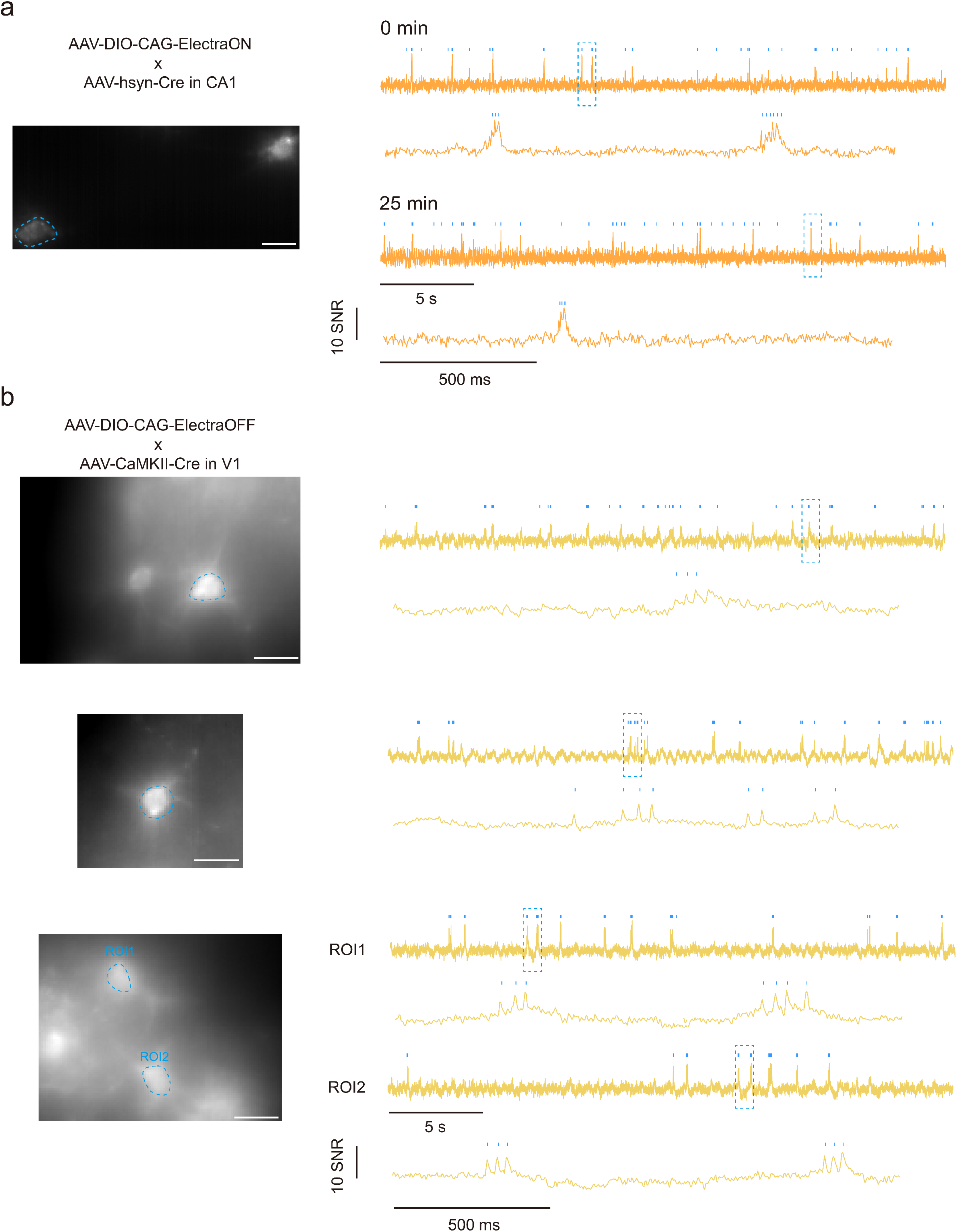
Voltage recording of ElectraOFF in the mouse brain using one-photon microscopy. **a.** Left, average projection image showing ElectraON-expressing neurons in mouse CA1 with the target neuronal ROI highlighted in blue. Right, 30-sec raw fluorescence trace of the neuron shown in the image at the beginning and at the 25th minute of recording. Trace snippets highlighted in dashed box were shown below. **b.** Same as a but for three FOVs showing L2/3 pyramidal neurons in V1 expressing ElectraOFF. Scale bar = 25μm.

